# Spatiotemporal Ecological Chaos Enables Gradual Evolutionary Diversification Without Niches or Tradeoffs

**DOI:** 10.1101/2022.05.25.493518

**Authors:** Aditya Mahadevan, Michael T. Pearce, Daniel S. Fisher

**Author notes:** Current institution: Meta Data Science (Michael T. Pearce). Authors contributed equally (Aditya Mahadevan, Michael T. Pearce).

## Abstract

Ecological and evolutionary dynamics are intrinsically entwined. On short time scales, ecological interactions determine the fate of new mutants and changes in the community they induce, while on longer time scales evolution shapes the whole community. How eco-evolutionary dynamics gives rise to the extensive coexisting diversity of strains found in many bacterial species is a major puzzle. In this paper we study the evolution of large numbers of closely related strains with generalized Lotka Volterra interactions but no niche structure. The host-pathogen-like interactions drive the ecological dynamics into a spatiotemporally chaotic state characterized by continual local blooms and busts. Upon the slow serial introduction of new strains, the community is found to diversify indefinitely, accommodating arbitrarily large numbers of strains in the absence of any kind of stabilizing niche interactions. This diversifying phase is robust to changes in evolutionary parameters, and persists even in the presence of a distribution of general, nonspecific fitness differences between individual strains, which explicitly break the assumption of tradeoffs inherent in much previous work. However, gradual increase of the general fitnesses in the ecosystem slows down the diversification. Quantitative analysis of the range of behaviors is carried out by a combination of analytical methods and simulations. Building on a dynamical-mean field-theory understanding of the ecological dynamics, an approximate effective model captures the effects of evolution on the distributions of key properties, such as strain abundances. This work establishes a potential scenario and a theoretical framework for understanding how the interplay between evolution and ecology can give rise to extensive fine-scale diversity. Future avenues for investigation are suggested, including the effects of the build-up of relatedness between strains, how conditioning on the evolutionary history affects the ecological interactions and dynamics, and application to coevolution of the diversity of a bacterial and a phage species.

## 1 Introduction

A remarkable discovery of the DNA sequencing revolution is the vast diversity of microbes [1, 2, 3, 4]. Increasingly it has become clear that this diversity extends far below the level of conventionally defined species to finer and finer genetic scales [5, 6, 7, 8], and in some species, a great multitude of strains coexist and compete in the same spatial location. Why doesn’t “survival of the fittest” drive almost all strains extinct, at least locally? Traditional explanations invoke the existence of a great many spatial or functional niches which limit competition between strains, down to “micro-niches” involving finer differences. But — especially for bacteria in relatively simple environments such as the marine cyanobacterium *Prochlorococcus* [6] — does it make sense to postulate nano- or pico-niches, *ad absurdum*? Or is a statistical description of the small subtle differences more appropriate? Community ecology models with many similar strains competing for a mixture of resources have been much studied, but in their simplest manifestations the maximum number of coexisting strains is limited by the number of resources — more generally, the number of chemicals via which they interact [9]. Perfect “tradeoffs” are sometimes invoked to enable higher diversity [10, 11, 12] but even tiny differences will destroy this [13].

An alternative to the multi-niche scenario is the *neutral theory* of ecology which postulates that species are similar enough that they are somehow ecologically equivalent, with their population dynamics dominated by stochastic births, deaths, and migration. The predictions of this theory for abundance and spatial distributions are intriguingly similar to some data [14, 15]. However for microbes with short generation times and huge populations without tight bottlenecks, the neutral scenario is not viable: Even if the differences between strains could be neglected over the long times for which they have coexisted, the dynamics from stochastic fluctuations are far too slow. Instead, rapid population dynamics with large changes of relative abundance are often observed [16, 17]. “Selection,” in the broad sense of differential population growth rates, is clearly involved. Thus if a highly diverse population *appears* “neutral” in some respects (including close-to-perfect tradeoffs) this must *emerge* from the complex ecological and evolutionary dynamics: It should not be assumed.

It is often said that pathogens promote diversity [18, 19, 20, 21]. But there is thus far little understanding of how or under what circumstances ongoing coevolution of hosts and pathogens could cause and sustain extensive coexisting within-species diversity. Understanding this process theoretically is a long-term goal, towards which the present work is a substantial step. To make progress, we need to distill the general phenomenon of fine-scale diversity to its most basic, and endeavor to develop potential scenarios in which evolution, coupled with ecology, might play out. For closely related strains, there is no compelling reason why interactions with siblings should be much stronger than those between distant cousins. Thus, we first ask: Without assuming niche-like interactions, perfect tradeoffs, or spatial gradients, can a highly diverse collection of closely related strains stably and robustly coexist? If so, can such a highly diverse “phase” evolve and continue to evolve and diversify? If the evolution is fast, some amount of diversity will always exist (although the common ancestors of the population at any time may be recent and few). Thus we consider the most difficult regime for diversity: when ecological and spatial population dynamics occur much faster than evolution.

In recent work [22] (referred to henceforth as PAF) we developed a new scenario for the coexistence of multiple closely related strains that are *assembled* all together into a community, leaving aside the question of their past or future evolution (or even how the community is assembled). In this scenario we explored a particular key feature of models of many similar strains: the nature of interactions between pairs of strains. Competition for resources in a well-mixed environment leads to positive correlations: if more *A* individuals are worse for *B*, then more *B* are worse for *A*. We consider the opposite case where the interactions are anticorrelated. This can arise if the competition is one-on-one: if *A* beats *B*, then *B* loses to *A*, so the effects on each other are opposite. More interesting biologically, is a spectrum of generalist phage strains that prey, with varying efficacies, on a spectrum of bacterial strains. If a particular phage strain, *a*, does better than average against a particular bacterial strain, *b*, then more *b* individuals are better for *a*, and more *a* are worse for *b*, leading to anticorrelated interactions.

Host-pathogen, and other anticorrelated interactions, give rise to “kill the winner” ecological dynamics [20]. If a strain rises to high abundance, other strains that do well against it will bloom and drive down the abundance of the first, and the process repeats. With many strains that do not have their own niches, this leads to wilder and wilder chaotic variations of abundances, soon driving most types extinct [PAF]. While we are particularly interested in coevolving bacteria-phage diversity, to build up an understanding of the complex eco-evolutionary dynamics, we focus in this paper on simpler models that — as we have shown in PAF — capture many of the key features.

The classical mechanism for maintaining diversity when fluctuations drive extinctions is to invoke a mainland, from which emigration of strains to the location (island) of interest can repopulate strains that have gone extinct [23]. However, this begs the question: How does the mainland evolve and sustain such extensive diversity? We showed that rudimentary spatial structure — a large set of *I* islands with a low migration rate between all pairs of islands — can maintain some of the diversity: Many strains go globally extinct, but a large fraction persists essentially indefinitely in a spatiotemporally chaotic phase (hereafter STC). On each island the abundances vary wildly and chaotically. But, crucially, the chaotic dynamics desynchronize across the islands allowing strains that go extinct locally to be repopulated from other islands. This mechanism is a manifestation of the “spatial storage effect” [24]. On each island, each persistent strain occasionally blooms up to high abundance and subsequently crashes. While it is at low abundance, dispersal from blooms on other islands rescues the strain from local extinction until conditions are favorable and its population blooms again, sends out migrants, and crashes. This STC is very robust: strains either go extinct rapidly, or persist globally for times that are exponentially long in the number of islands.

Complementary work [25, 26] suggests the generality of the STC beyond anticorrelated pairwise interactions, although in the Lotka-Volterra models studied in these works of Roy et al., the diversity is limited by the strength of niche (or self-) interactions, which also limits the diversity of stable communities (see Section 2.1.1). Indeed, much previous work has focused on ecological dynamics that reach a stable state, where diversity is limited by strength of niche interactions compared to inter-species interactions [27]. Here we approach evolutionary dynamics in similar generalized Lotka-Volterra models, but from the opposite starting point: all interactions are of comparable magnitude which makes the effects of self-interactions negligible compared to the effects of the total interactions from all other strains. Then there is no large stable community, and spatial structure together with chaotic dynamics is needed to maintain diversity.

With anticorrelated interactions, arbitrarily large numbers of strains can coexist in the STC even when spatial mixing — and hence competition — occur on timescales comparable to those of the local ecological dynamics. However if the strains differ somewhat in their overall growth rate, or other ways that make some generally better, this can limit the diversity. A natural assumption is that, having all survived on evolutionary time scales, the persistent strains will be similar enough that such differences are very small — but this assumption should surely be questioned: doing so is one of the goals of this paper.

Many theoretical (and some experimental) analyses, have, like our prior work, focused on ecological communities that are assembled without conditioning on their evolutionary histories: a number of species (or strains) is brought together, and the resulting community consists of the species that do not go extinct [27, 28, 29, 30, 31, 32]. Although this is an important starting point, it is essential to incorporate evolution to understand how the processes of mutation, inheritance, selection, and extinction could give rise to highly diverse communities.

Previous theoretical work has shown that diverse communities in certain consumer-resource models are destabilized by evolution [33], at odds with the highly diverse continuously evolving microbial populations in Nature. Others have focused on eco-evolutionary dynamics when the mutation rate is high enough to sustain diversity: but in this case the common ancestor of coexisting strains is recent since the balance between mutation and extinction is responsible for maintaining diversity [34]. Therefore extensive diversity over a wide range of genetic differences does not have time to evolve. Yet others have shown that when niches in phenotype space are assumed, boom-bust dynamics can result in the evolution of higher diversity than stable equilibrium dynamics [35]. But overall there is no clear consensus on whether evolution tends to destabilize or to increase diversity in ecologically interacting communities — indeed the answer to this question is likely context-dependent — though observations of the natural world suggest that evolution often results in increased diversity. Here we investigate evolution starting from a state with spatiotemporally chaotic ecological dynamics as studied in PAF, where niches are absent and the diversity — at least initially — is stabilized by the dynamic ecological fluctuations.

We are interested in understanding diversity that has existed for a very broad spectrum of evolutionary time scales, far longer than ecological or spatial mixing time scales. We thus study the extreme limit where the mutation rate is small enough that the ecological and migratory dynamics reach steady state before the introduction of each new strain. This *quasistatic* limit of evolution is the “hardest” for diversification. In addition, and in contrast to some previous work [36], we assume that global extinctions of a strain are permanent: an extinct organism cannot be resurrected even if conditions later become favorable for it. Focusing on the STC phase, we endeavor to answer: Can a highly diverse STC phase evolve? Under what conditions? Can this phase continue to diversify? Is the diversity stable to *general fitness* mutations that are not artificially constrained by assumptions of tradeoffs?

The structure of this paper is as follows: Section 2 introduces the main model and its relation to previous work. Section 3 describes the phenomenology of an evolving community in the STC phase, looking at the effects of correlated mutants, interaction statistics, and general mutations on the ecological diversification. Then Section 4 develops the theory and analysis that are needed to understand these phenomena. Building upon the dynamical mean field theory developed in PAF, we can understand the criterion for invasion and survival of a new strain simply in terms of emergent quantities. We present an approximate framework for understanding the evolutionary dynamics in our model, and compare it with simulations. Finally Section 5 raises additional questions and discusses possible extensions.

## 2 Models

We now define the model, building up from local deterministic population dynamics, to spatial migration and demographic stochasticity, to evolutionary dynamics.

### 2.1 Ecological interactions and dynamics

We consider a community initially of *K* strains with all possible pairwise interactions between them. A paradigmatic model is the generalized Lotka-Volterra model [37], with strain *i* having an intrinsic population growth rate *r*_*i*_, modulated by the effect of each strain *j* on strain *i* parametrized by the interaction matrix element *W*_*ij*_. Denoting the population of strain *i* by *n*_*i*_, one has 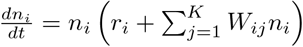 for each of the *K* strains, with competitive interactions corresponding to negative *W*_*ij*_. It is by now traditional to draw the values *W*_*ij*_ from a random distribution, with the hope that some results from such models will be valid for classes of complex ecologies, albeit not about specific systems [38]. Traditionally, the effect of a particular strain on itself, *W*_*ii*_ < 0, is treated as special — assuming each strain has its own niche — and interactions with the others (niche overlap) are assumed much smaller in magnitude. However, we are interested in closely related strains for which the interactions are all quite similar so the variations in the *W*_*ij*_ are small compared to their negative mean, *Ŵ* < 0. Likewise the *r*_*i*_ vary slightly around some mean 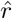. The total population will then be roughly fixed at some *N*, by the balance between the effects of 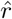 and negative *Ŵ*. It is convenient to replace these large terms by a Lagrange multiplier ϒ(*t*) that fixes the total population to Σ_*i*_ *n*_*i*_ = *N*. and change to working with fractional abundances, *ν*_*i*_ = *n*_*i*_/*N*. This parameterization is equivalent to the oft-studied replicator equations [39, 40, 41].

The deviations from average of the intrinsic growth rate of a strain together with the total interaction on it can be combined to yield *general, nonspecific fitness differences* between the strains: 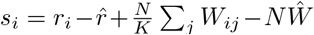 — emphasizing that their average interactions with others can give a nonspecific advantage or disadvantage to different strains relative to the full pool of strains. We will refer, loosely, to the *s*_*i*_ of a strain as its *general fitness* and take it to be *independent of the community* of strains with which it interacts. The residual interactions between strains, arising from small deviations from the average interactions, can be written as 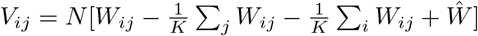. In terms of the {*ν*_*i*_}, {*V*_*ij*_} and {*s*_*i*_}, the Lagrange multiplier takes the form 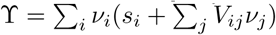.

Since the *V*_*ij*_ and *s*_*i*_ are sums and differences of similar magnitude terms, it is natural to approximate them as random variables with the hope that the model will yield behaviors that are robust to specific choices of their statistics: testing this assumption is one of the goals of this paper. For simplicity we choose E[*V*_*ij*_] = 0, and 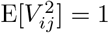 for *i* ≠ *j* — setting the overall ecological time scales — and the covariances zero except for correlations between how *i* and *j* affect each other with E[*V*_*ij*_*V*_*ji*_] = *γ*; (for convenience we choose 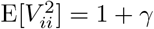 but this has negligible effect in large communities).

The crucial parameter, *γ*, controls whether the interactions are mainly competitive (*γ* > 0) or host-pathogen like (*γ* < 0), on which we focus. We have shown that random interaction matrices with such anticorrelations behave very similarly to host-pathogen models with the appropriate block sub-matrix structure, as discussed further in Section 5.4.

[Note that our parameterization of the generalized Lotka-Volterra model is different from the parameterization commonly seen in the literature [25, 27] in that it does not rescale the interaction strengths by the number of strains, which is a mathematically convenient (but biologically unmotivated) choice that allows one to take the infinite-strain limit while maintaining finite total interaction strength. It is more natural is to have *K* a free parameter, especially for studying evolution, where new strains can be added and others go extinct — so the number of strains in the community will vary on evolutionary time scales.]

#### 2.1.1 Stable communities and niche models

Much of the theoretical work on complex communities has focused on the interplay between strong niche interactions and much weaker and roughly random inter-species interactions. We define the niche-interaction parameter *Q* = − E[*V*_*ii*_]: strongly positive *Q* (relative to the other *V*_*ij*_ which have variance unity) results in a unique stable community with a substantial fraction (≥ 1/2) of the *K* assembled strains surviving. However for *Q* ≫ 1, and no general fitness differences, the community becomes unstable when 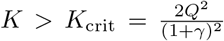, at which point precisely 1/2 of the *K* assembled strains survive [42, 43, 44]. For *K* > *K*_crit_, complicated dynamics ensue which are only very partially understood [45, 25].

Throughout this paper we set *Q* = 0, so that there is no large stable uninvadable community. Note that if *Q* were positive but not large, then its effects would be small as the population of each strain is typically of order 1/*K*. There is a special case of perfectly antisymmetric *V*_*ij*_ (*γ* = −1 with *Q* = 0) — here referred to as the ASM — which is exactly marginal with very special properties analyzed in PAF: its behavior underlies much of the understanding of the −1 < *γ* < 0 models of primary interest here.

### 2.2 Spatial dynamics

We study the simplest model with spatial structure: a large number, *I*, of identical islands (or demes) with interactions only within each island and migration between pairs of them. With Greek indices labeling islands the dynamics of the abundance are given by

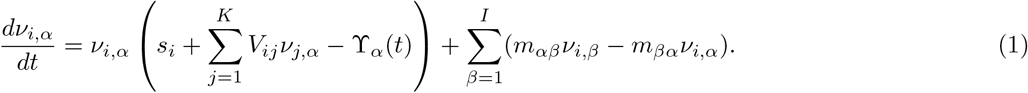

with *m*_*αβ*_ the migration rate (per individual) from island *β* to island *α* and the local Lagrange multiplier, ϒ_*α*_, keeping the total population on island *α* fixed at *N*, (i.e. Σ_*i*_ *ν*_*i,α*_ = 1 for each island). Here, we focus on the spatial mean field limit in which the migration rate is the same, *m*/*I*, between every pair of islands. The total migration of strain *i* into and out of island *α* is then simply 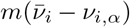 with 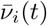 the average across islands. The magnitude, *m* of the migration rate, is of fundamental importance. If *m* is too small the migration is too rare to repopulate islands after local extinctions. If *m* is too large and the local dynamics is chaotic, the chaos will synchronize across the islands and the total population of each strain will fluctuate wildly, rapidly driving most strains extinct. We will focus on the wide intermediate *m* regime, which spans several orders of magnitude for large *K* [PAF].

### 2.3 Demographic fluctuations, extinctions, and recolonization

With large populations on each island, the demographic fluctuations have little effect on the dynamics. Even when the local population of a strain is small, if it has positive growth rate fluctuations will not much matter, while if it has negative growth rate it will go deterministically extinct. The extinction threshold in Figure 1 is indicated by the horizontal purple line: its value does not much affect the behavior as long as it is much below lower limit of frequency stabilized by migration — which we term the *migration floor*: *ν*_floor_ which is proportional to 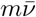, the migration rate of a strain into the island. For strains near local extinction (when the fractional abundance drops below 1/*N*) demographic fluctuations are potentially important. But with 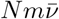 very large, local extinctions will be rare: this is the mostly-deterministic regime on which we focus. Nevertheless, for some strains the total population will decrease systematically (albeit possibly with temporary fluctuations up caused by local blooms) and, for convenience, we set a global extinction threshold of 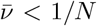 with the choice of *N* not mattering much as long as *mN* ≫ 1, to which we restrict consideration. In Section 5.3 we comment on the effects of local extinctions in the context of real spatial dynamics.

**Figure 1:**
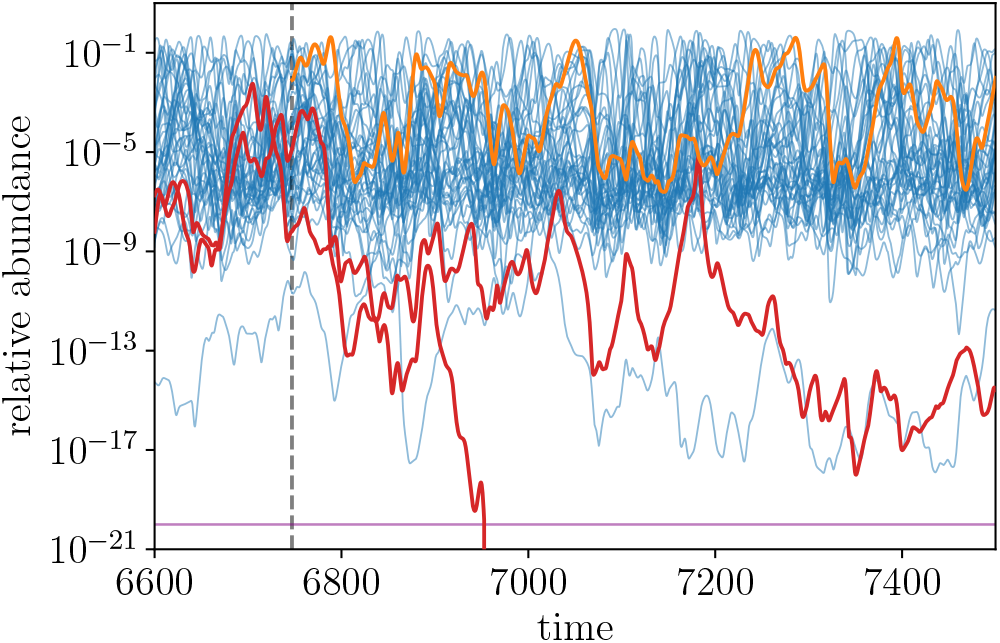
Chaotic dynamics of strain abundances on a single island in the spatiotemporally chaotic phase (STC). (For clarity, only one third of the *L* = 125 persistent strains are shown.) Each persistent strain occasionally blooms up to high abundance and between blooms its abundance is sustained above a migration floor (here ≅ 10^−9^) set by migration from other islands, though a few marginal strains fluctuate below this threshold. At the beginning of a new *epoch* (vertical dashed line), a new independent strain, colored in orange, is introduced: in this case, it fares well enough on average to persist. The interactions of the new strain with the members of the community cause the two red strains, which persisted in the previous epoch, to go globally extinct (one rapidly and the other before the end of the epoch, only a portion of which is shown).

### 2.4 Evolutionary dynamics

Our primary focus is the effect of evolution on communities. We consider rare invasions either of mutants that are correlated with their parents, or of unrelated invaders. Each added strain, labeled *A*, is parameterized by its interactions, *V*_*Aj*_ and *V*_*jA*_ with all the other strains present in the community and its general fitness, *s*_*A*_. For unrelated invaders, the new *V* and *s* are drawn from the same distributions as the original strains — which have evolved and whose statistics therefore differ from the original strain statistics, the extent of which will be explored.

The general fitnesses, *s*_*i*_, play an important role — especially during evolution. A natural assumption is that well adapted strains that have all survived will have similar *s*_*i*_ and thus a narrow *s*_*i*_ distribution. But we must show the self-consistency or not of this assumption: thus we consider several classes of the distribution of the assumed-independent *s*_*i*_.

We consider a family of distributions of *s* of the form:

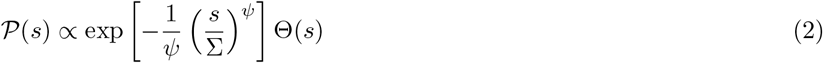

with characteristic scale Σ: this form is motivated by the expectation that the tail of the *s* distribution is particularly important for evolution, and the Heaviside function Θ(*s*) constrains the general fitnesses to be positive, though in practice it is *s*_*i*_ relative to, *ŝ*, the *mean general fitness of the extant population* that matters for the fate of strain *i*, as will be discussed.

For adding mutants, a parent, labeled *P*, is chosen with probability proportional to its mean abundance 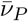 and a mutant daughter, labeled *M*, is drawn with interactions satisfying Corr[*V*_*kM*_, *V*_*kP*_] = Corr[*V*_*Mk*_*, V*_*P* *k*_] = *ρ* for all strains *k*. The details of this scheme for *k* = *M, P* are discussed in Section 6.3. Similarly, *s*_*M*_ is chosen to be correlated with *s*_*P*_ as detailed in Section 3.6.

We conduct simulations in *epochs* which are run long enough that the ecological and migratory dynamics have reached a steady state, with some fraction of the strains having gone *permanently extinct* globally, leaving *L* persistent strains. A single new strain is then introduced and the process repeated. The actual process of invasion from low abundance on one island is complicated, and often leads to failure. To avoid a proliferation of such failed runs, we ask instead whether the invader *could* successfully invade and persist if it were lucky initially. Thus we set the mutant’s abundance to 1/*L* on all the islands at the same time (and proportionately decrease the abundances of the other strains to maintain Σ_*i*_ *ν*_*i,α*_ = 1).

### 2.5 Timescales

There are multiple time scales involved in the dynamics. The basic time scale for *differential growth or decay* of populations of strains is primarily set by the magnitude of the interactions and the number, *L*, of extant strains. If they each have abundances of order 1/*L*, the total interaction on each is roughly the sum of L random terms each of order *V*/*L*: with the *V* chosen to have variance unity, the corresponding time scale is 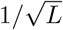: this is indeed the time scale over which the persistent effects of interactions on a strain are effective. When, in addition, there are general fitness differences, these will have substantial effects in time 1/*σ*_*s*_ with *σ*_*s*_ roughly the width of the *s*_*i*_ distribution of the *extant* strains.

Each strain’s local abundance can vary over a wide logarithmic range. In the STC phase, abundances range from high abundance down to the migration floor set by migration from other islands. This logarithmic range is approximately *M* ≡ log(1/*m*) which is quite large for the typical *m* = 10^−5^ that we use in simulations. The time for abundances to fluctuate from large to small — the *duration of blooms* — is 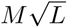. The time to reach equilibration is determined by the strains that are just barely going extinct. We show that in evolved communities this is longer by an extra factor of 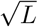: thus the *ecological equilibration time* for a community is of order *ML*.

Although it will not play much role here, there is also a short time scale associated with the dynamic fluctuations: roughly the time that a strain spends near the peak of a bloom, which is similar to the inverse of its instantaneous growth or decay rate. This is dominated by the net effect of its interactions with the small subset of 𝒪(*L*/*M*) strains that happen to be abundant at that time: the variance of this is 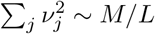, which makes the *dynamic fluctuation time* ~ 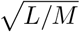. Between this dynamic fluctuation time scale and the time scale for blooms, correlations in the growth rates decay as a power of the time difference. During a bloom, each strain experiences multiple reversals from growth to decay: this is a special property of the self-organized chaotic state.

The time to go extinct for a strain destined to do so depends logarithmically on the extinction threshold 1/*N*, but as long as *Nm* is very large, whether extinctions occur is not strongly dependent on *N*, as analyzed in PAF. In our simulations we choose for convenience *Nm* ≫ 1 so that log(*N*) is a few times log(1/*m*).

The time scale for migration to be effective would, if there were no differences between the strains, be of order 1/*m* which is very long. However, the spatial dynamics are much faster than this because of the exponential growth of local populations when they happen to be in a favorable community. This makes the time scale for *exponential spread across islands* of a successful invader be on the order of the bloom time on a single island, 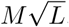. This is analogous to the rapid spread of a Fisher wave driven by selection, even when the spatial dynamics is diffusive [46, 47].

The time scale of demographic fluctuations — even if some strains were phenotypically identical — would be very slow ~ *Nτ*_*gen*_ with *τ*_*gen*_ ≪ 1 (in our units) a generation time. In practice, these fluctuations only matter when the populations happens to be very small and being driven extinct on the deterministic dynamic time scale ~ 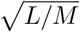. But as long as 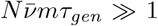, there are many migrants arriving and the dynamics is essentially deterministic. If the island-average 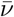 drops enough then the migrations become stochastic with time intervals between them of order 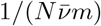. But when this occurs for substantial time, the global population is likely to be on the way to extinction. We focus on *Nm* ≫ *L* for which the population dynamics of the persistent strains are not strongly influenced by the stochastic or migratory demographic fluctuations.

We have chosen the *evolutionary time scale* to be much longer than all the other important time scales. Thus each epoch before a new mutant or invader is added is chosen to be several times the ecological equilibration time, typically 3*ML*. We show in Section 6.2 that increasing this epoch length by a factor of 10 makes little difference. If the time between adding new strains were much longer than this, evolution would depend somewhat on the number of islands. While the typical times for persistent strains to fluctuate to extinction — by unluckily sustained bad conditions on many islands — is exponentially long in the number of islands, for close-to-marginal strains, fluctuations to extinction are not so rare. But, as discussed later, the extinction of these strains would not much change the evolutionary behavior. (Note, however, that as *ρ* becomes close to 1, the timescale for the very-similar parents and mutants to potentially drive each other extinct diverges, so for this one must consider longer epochs as in Figure 10.)

## 3 Phenomenology

The key properties of the STC phase, as summarized above, are chaotic coexistence of all the strains on each island, with the local abundances fluctuating over a range in log *ν* of *M* ≡ log(1/*m*). Crucially, the chaos desynchronizes across the islands. Some strains go globally extinct but each persistent strain on each island occasionally has a bloom up to high abundance (*ν* ~ *M*/*L*). The emigr/ants from these blooms sustain the population of the strain on the other islands above the migration floor of 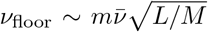 (the last factor being the inverse of a typical instantaneous population decay rate discussed above). We chose large enough population size that the island population at the migration floor, *Nν*_floor_, is very large for all strains that are not going globally extinct. A snapshot of the abundances on each island shows the strains distributed roughly uniformly in log(*ν*) down to the migration floor, only occasionally fluctuating substantially lower (Figure 1).

As shown in PAF, the STC occurs for a wide range of migration rates, and for a range of negative values of the symmetry parameter, *γ*. It is very robust, with the strains persisting for times that are exponentially long in the number of islands. However some of the strains are close to marginal and can be driven extinct by not-so-rare fluctuations even with the *I* = 40 islands we study. These can also be driven extinct by perturbations such as an invading strain: a process that we need to analyze in detail.

Each strain in the STC phase (and more generally in other types of steady state communities) can be characterized by its *local* invasion eigenvalue, which we call its *bias*: this is the average rate of growth (or decay) of a strain’s population on a single island when there is no migration and the strain has low enough abundance that it does not affect (and has not affected in the recent past) the other strains. The bias can be written as *ξ*_*i*_ = *s*_*i*_ + Σ_*j*_ *V*_*ij*_ ⟨*ν*_*j*\*i*_⟩ − ⟨ϒ_\*i*_⟩ where the notation ⟨*ν*_*j*\*i*_⟩ and ⟨ϒ_\*i*_⟩ denote the time-averaged abundance of strain *j* and average Lagrange multiplier *in the absence of strain i*. (Note that ϒ is not much changed by the absence of the one strain, but the abundances of the others, collectively, are in small but essential ways affected by whether or not the strain of interest is present, as discussed in Section 4.1.)

With many strains participating in the chaos on each island, and desynchronization across islands, we expect the chaos to be ergodic, so that the time averages and spatial (across islands) averages of all quantities are equal in the steady state. Therefore we can use the notation 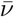 (for spatial average) instead of the time-average notation ⟨*ν*⟩ (and similarly for other quantities). We will use the overbar notation, except when conceptually the time average is clearer. In practice, our *I* = 40 islands are enough that the persistence times of almost all surviving strains are very long and averages across islands of the more important quantities do not fluctuate much in steady state.

The biases of the strains are of fundamental importance in characterizing their behavior and the effects of evolution on the ecology. Each strain has its own bias, which determines, in a complicated way, its average abundance 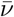. An important characteristic of a collection of strains in ecological steady-state is the *distribution* of their biases. Strains with too-negative bias go globally extinct. But a crucial feature of the STC phase is that the desynchronization across islands allows strains with somewhat negative bias to persist forever (in the infinite island limit). While many of these persistent strains spend most of their time at low abundance, rare blooms to high abundance of each persistent strain are always occurring on a finite fraction of the islands and migration from these prevents the strain from going globally extinct. This stabilization of some negative-bias strains is enabled by a feature of the STC phase: the timescale of the blooms is the same as the time scales set by the systematic changes in log(*ν*_*i*_) caused by the bias — this property of the endogenous fluctuations keeps the system tuned to the special self-consistent chaotic state [22].

Despite the possibility of rescue from extinction via rare blooms, there is a critical negative bias (sharp for large *L* and large *I*) below which strains no longer persist even in the infinite island limit. For strains with *ξ* below this negative critical bias *ξ*_*c*_ (which depends on the parameters and the number of strains), blooms up to high abundance are not frequent enough to repopulate islands that have experienced local extinctions (or even dips far below the average migration floor) and deterministic global extinction of such strains ensues. In the limit of large *I* and large *L*, strains with *ξ* < *ξ*_*c*_ go extinct, while strains with *ξ* > *ξ*_*c*_ persist indefinitely. Finite *L* and finite *I* effects, together with the finite time for each epoch, will round out the sharpness of the borderline between persistent and extinct. However the marginal strains involved have little effect on others and whether or not they persist does not much matter for the current epoch.

### 3.1 Continual assembly and diversification

The evolutionary process we study starts from an assembled collection of *K* unrelated strains. After the ecological and migratory dynamics have reached steady state, some of the strains will persist: we call the size of this persistent community *L*_0_. The *K* − *L*_0_ strains that have gone globally extinct are permanently removed (in contrast to mainland models).

When a new strain is introduced into the ecosystem, if it successfully invades it perturbs the biases of the extant strains, and can trigger extinctions of some of them by shifting their bias to below *ξ*_*c*_ (Figure 1). We study the slowly evolving regime in which the ecosystem dynamics reach steady state between each introduction of a new strain. The number of persistent strains, as a function of the number of invasion attempts, *Y*, we denote by *L*(*Y*): this is of fundamental interest.

We first describe the evolutionary dynamics when the general fitness differences between the strains, {*s*_*i*_}, can be neglected. The simplest to understand is if the invaders are unrelated: i.e. the interactions that characterize them are independent of those of the extant (or already-extinct) strains.

For *γ* = −0.8 and unrelated invaders (*ρ* = 0), multiple runs starting with different sets of *K* = 50 initial strains reveal that around one fifth of the replicates enter a steadily diversifying regime in which *L* increases roughly linearly with the number of attempted invasions, at a rate of around 0.25 per attempt. The other replicates crash under the disturbances of the invasions down to only a few persistent strains. Subsequent invasions can cause *L*(*Y*) to increase somewhat, but it quickly crashes back down and the community does not steadily diversify (Figure 2A). This low diversity regime that occurs after a crash is discussed further in Section 4.9 and Figure 13.

**Figure 2:**
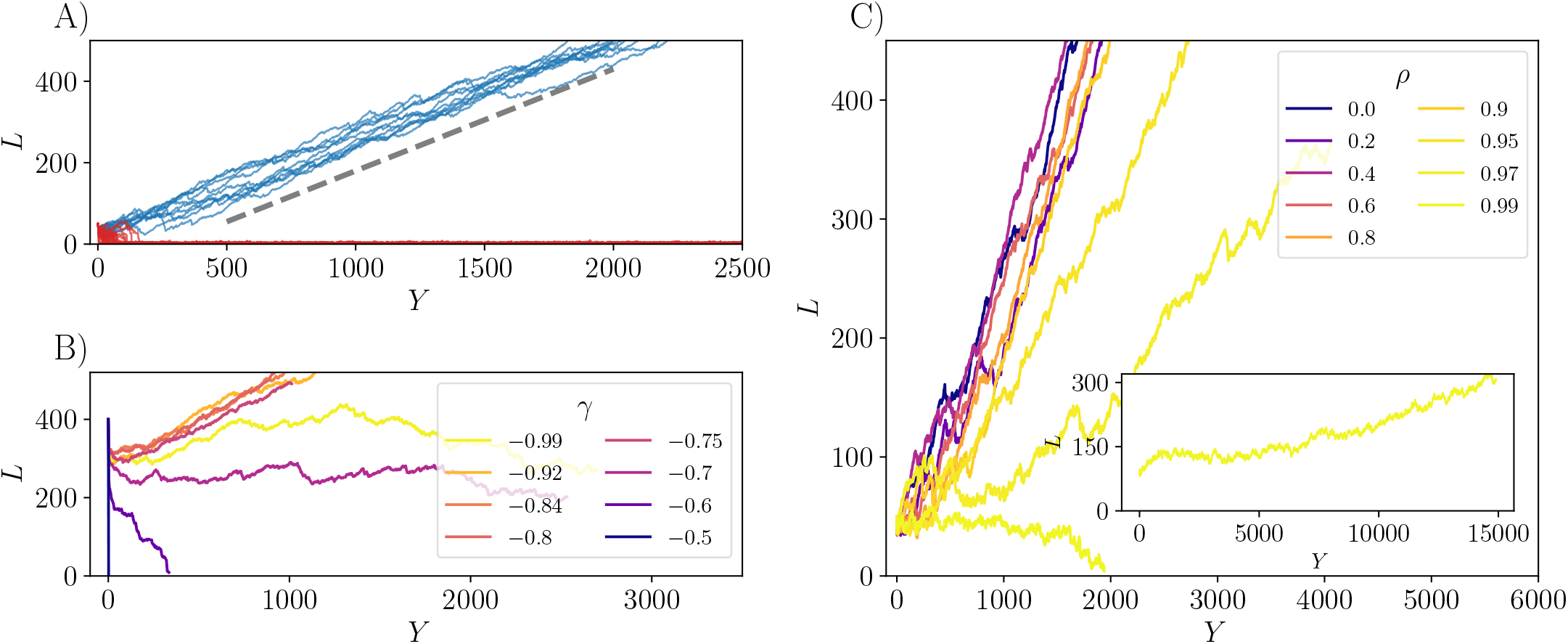
Evolution of number of strains without general fitness differences. (A) With *γ* = −0.8, *m* = 10^−5^, and initial number of strains *K* = 50, most initial communities (red) crash and fail to recover, while others (about 20%, blue) continually diversify. Quantities are shown versus the number of attempted invasions, *Y*. Once the communities are large, around 80% of invasions are successful and the mean number of extinctions per successful invasion is ≅ 0.7 (Figure 3) so that on average the number of strains in the community grows linearly by *U* ≅ 0.25 per invasion attempt (dashed line). (B) Whether diversification occurs, and its rate if it does, depends on the symmetry parameter, *γ*. There are two transitions as *γ* becomes less negative (seen here with *K* = 400). For any *γ* ≳ −0.7, introduction of new strains reduces the diversity. For even less negative *γ*, the STC breaks down and the diversity crashes immediately. For more negative *γ*, steady diversification occurs, fastest here for *γ* ≳ −0.8, though again slowing down as *γ* → −1. (C) Evolving communities under successive introduction of *mutants*, each with correlation *ρ* with its parent (here *γ* = −0.8). The diversification rate varies nonmonotonically with *ρ*, with fastest diversification for *ρ* ≈ 0.8. There is a significant slowdown for *ρ* close to unity — small effect mutations. Here *K* = 50 but only trajectories that do not crash are shown. With very small effect mutants, *ρ* = 0.99, the evolving community is very susceptible to crashing from *K* = 50, however the inset shows that it is possible to reach a regime of steady diversification starting from a larger *K* = 100.

The observations with particular parameters illustrate one of the crucial findings of this work: spatiotemporally chaotic ecological dynamics can allow — but do not guarantee — gradual strain-level diversification up to arbitrarily high number of strains. The behavior depends on the symmetry parameter *γ*, which must be substantially negative for the STC to exist. The simulations, Figure 2B, show that the average rate of diversification, *U* ≡ *d*⟨*L*/*dY*⟩, is nonmonotonic in *γ*, with slow diversification close to *γ* = −1 (its lower limit). As *γ* becomes substantially less negative, the STC still supports chaotic coexistence of many strains (illustrated here by *L*_0_ still large), but the system is unstable to the introduction of multiple new strains, with the diversity steadily decreasing, or quickly crashing, triggered by the evolutionary dynamics. The community diversifies most rapidly for *γ* ≈ −0.8. As we are interested in what can happen with various other additional features, we chose *γ* = −0.8 and *m* = 10^−5^ for all further simulations as shorter runs are needed near these values. We expect that the qualitative conclusions will be similar for a range of *γ* and *m* around these.

### 3.2 Evolution with correlated mutants

We next study proper evolution i.e. via mutations of existing strains. At the start of each epoch, a parent to mutate is chosen at random with probability proportional to its mean abundance. The interactions of the mutant with other strains are drawn from the same distribution as the original set, but with correlation *ρ* with the interactions of the parent. (The direct interactions between the parent and mutant have to be chosen separately as specified in Section 6.3 but, as they only account for a small fraction of the total abundance in diverse communities, the specific choice is not important.) As a function of *ρ* (with *γ* = −0.8), the rate of diversification is nonmonotonic being fastest for *ρ* ≈ 0.8, and only weakly varying for smaller *ρ* (Figure 2C). As *ρ* nears 1, the mutant and parent are more similar, and it becomes harder for them to coexist, since any difference between them will result in a systematic change in their relative abundance, eventually driving one of them to extinction (Section 4.6). Since *L* can only increase when both the mutant and parent coexist, increasing *ρ* eventually slows the rate of diversification. However this only occurs for *ρ* very close to unity and at this point we do not understand why such small effect mutations are needed before the rate of diversification is substantially suppressed. Of particular relevance is understanding what sets the probability of a mutant replacing its parent, given that it invades successfully. One might expect this to be close to one for high *ρ*, but it appears to be so only for *ρ* very near unity.

One complication that occurs as *ρ* increases towards 1 is that there is a slow transient in both the ecological and evolutionary dynamics. The system can only reach a steadily diversifying state when all the extant strains have been generated by the same mutational process that acts to generate the new mutants. As the correlation between a *g*^th^ generation offspring and its ancestor is of order *ρ*^*g*^, it takes at least 1/(1 − *ρ*) successful descendants to produce two relatively uncorrelated mutants from the same parental lineage — therefore it takes at least *L*_0_/(1 − *ρ*) successful invasions before the ecosystem reaches a steady state with *L* growing steadily. For *ρ* = 0.99, the simulations are just starting to reach this regime in the inset of Figure 2. However we suspect that even if *L*(*Y*) does eventually grow linearly, it will do so at a much smaller rate for *ρ* even closer to 1. We further discuss the parental replacement probability in Section 4.6.

### 3.3 Extinctions, diversity crashes, and nucleation from low diversity

A crucial question is whether it is possible to build up a highly diverse community from a small initial number of strains and, if so, on what this depends. Starting simulations at values of *K* between 10 and 90, we observe a crossover size, *K*^*^, of the initial number of strains above which diversification is robust, and below which the diversity typically crashes under the evolutionary dynamics and does not again increase substantially for the duration of the simulations, (Figure 3A). The presence of a crossover *K*^*^ suggests that once the system is sufficiently diverse, it tends toward further diversity. Thus the main obstacle to diversification in these models is going from a single strain to ≅ 50 strains. We define *K*^*^ heuristically as the lowest *K* that diversifies with probability more than 1/2 (see Figure 16). For each *K* there is a corresponding *L*_0_(*K*): the number of strains that persists after the initial drop in diversity. This somewhat-variable *L*_0_ appears to be the primary determinant of a community’s stability to evolutionary perturbations. However this stability also depends on the community’s history. For example, if the community has evolved gradually from a smaller number of strains to some current size *L*_*E*_, its response to continued evolution may be different than if it has been assembled from an initial number *K* that dropped through its initial evolution to an ecologically stable community of the same size, *L*_*E*_. This motivates definition of a crossover size *L*^*^ which is somewhat different than *L*_0_(*K*^*^), and is given by the minimum size of an *evolved* community for which the probability of crossing into the diversification regime is greater than 1/2 (see Section 4.9). We discuss the differences between assembled and gradually evolved communities further in Section 4.8, but leave an analysis of how these properties affect the nucleation probability for future work. The size *L*^*^ is a natural quantity to work with, as it is the relevant one when considering how a community might nucleate from a small number of strains into a diversifying regime. As we discuss later, we expect *L*^*^ < *L*_0_(*K*^*^) since just-assembled communities are more susceptible to perturbations than gradually assembled ones.

**Figure 3:**
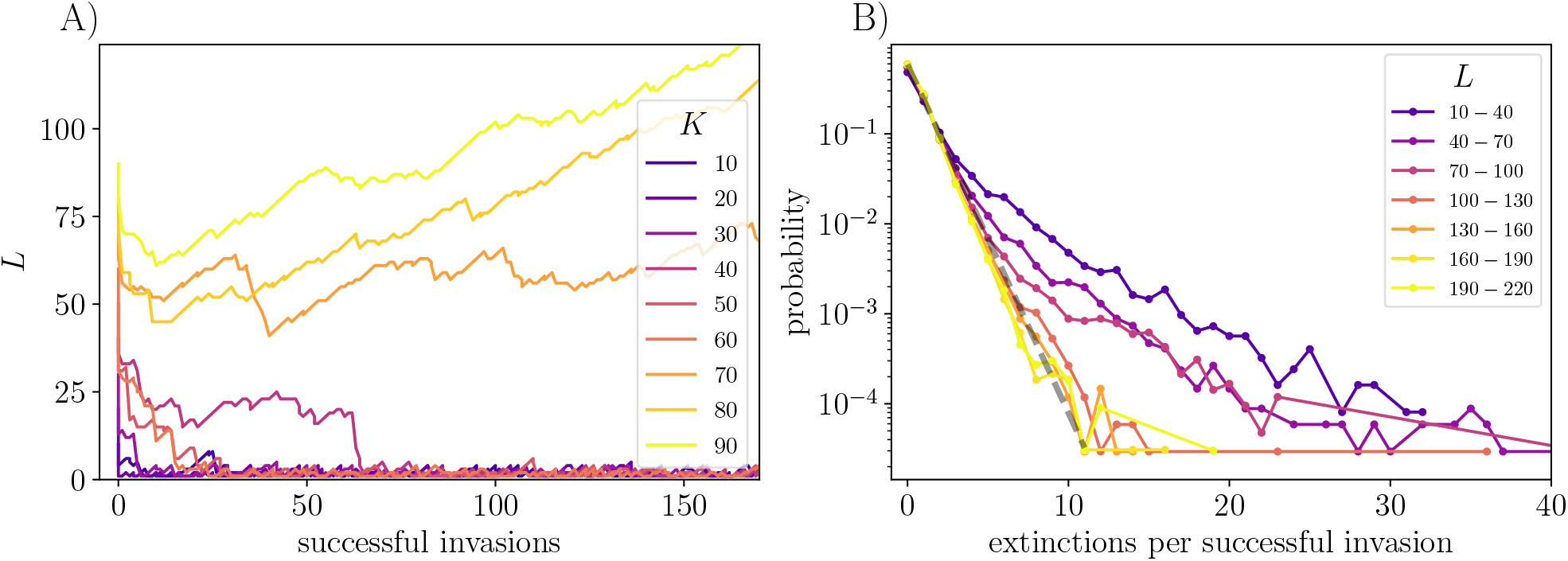
Crashes and distribution of number of extinctions per invasion. (A) The probability of a community crashing depends on the number, *K*, of initial strains. The characteristic size below which crashes dominate is ≅ 50 for the parameters *γ* = −0.8 and *ρ* = 0 as shown here. (B) The distribution of the number of extinctions per successful invasion depends on the size of the community. Data are aggregated from a range of *K*’s, each run with 100 replicates. For small community size, *L*, invasions occur in which a substantial fraction of the strains go extinct. But for large *L*, multiple extinction events are very rare and the distribution is close to geometric: the dotted line is 𝒫(𝓁) = (1 − *α*)*α*^𝓁^ for the probability of 𝓁 extinctions, with *α* ≅ 0.41, corresponding to an average of 0.7 extinctions per successful invasion as shown in Figure 2A.

The crossover size, *L*^*^, above which diversification is likely, is a function of *m, γ* and *ρ*, though we have not explored this in detail. For some range of *γ* (Figure 2B), *L*^*^ is infinite: the system always eventually crashes. The role of *m* in determining *L*^*^ can be partially understood from the mechanism of diversity crashes. One way in which these occur is when the STC disappears due to synchronization of dynamics between islands. This occurs more frequently when *m* is larger, but its dynamics are subtle because of local blooms up to high abundance which can can have outsize effects. Though we have not studied the synchronization and diversity crash process here, it suggests an interpretation of our simulation results (Figure 16) which show that decreasing *m* lowers *K*^*^, presumably because it makes the the STC more robust to synchronization.

We can quantify how likely the system is to diversify or crash by inspecting the number of extinctions, 𝓁, that occur for each invader that successfully enters the ecosystem. The distribution of 𝓁 is dependent on *L* and shows that more diverse ecosystems suffer on average fewer extinctions triggered by invasion of each new strain (Figure 3B). For small communities *L* of order *L*^*^ or less, extinctions of a substantial fraction of the strains occur, and cascades of extinctions in response to successive invasions cause *L* to crash. The fragility of the communities with *L* ≅ *L*^*^ to evolutionary perturbations and their strong tendency not to recover after a crash (Figure 2A), implies that the process by which a low diversity community *could* diversify is very different than the steady diversification of already large communities. The transition from the low diversity to the diversifying regimes must be mediated by a very rare nucleation event in which — roughly — *L* becomes larger than *L*^*^. We discuss the low diversity regime and speculate about such nucleations in Section 4.9. In the steadily diversifying regime the distribution of extinctions per successful invasion is close to exponential — a result that we derive in Section 4.3. For large *L*, the chances that a substantial fraction of the strains go extinct is extremely small and decreases exponentially as *L* increases further: thus for large *L* the continual diversification is essentially deterministic.

### 3.4 Evolutionary dynamics with general fitness differences

So far we have observed that when mutants or invaders differ only by their interactions with each other, there is robust and rapid diversification, provided that the initial diversity in the STC phase is high enough. Since we have not enforced any precise constraints or perfect tradeoffs, it is possible for a strain to be better than average against the others in a general way: this is equivalent to having a small general advantage, larger *s*. If strains have such an advantage they will be more likely to persist both ecologically, and evolutionarily. In the community that exists after the initial extinctions f rom a randomly assembled collection of strains, a persistent strain, say *i*, will typically already have better than average Σ_*j*_ *V*_*ij*_/*L*_0_. However this will be atypically large by an amount only of order 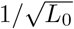. It is possible that under evolution, the average interactions that persistent strains feel continue to increase (Section 4.8). However with unrelated invaders the observation of steadily increasing *L*(*Y*), implies this cannot occur: such strains are too rare to be “found”. With mutants, rather than unrelated invaders, the situation could be different but our simulations show no sign of evolution of such anomalously good strains.

We now study the effects of *explicitly including general fitness differences*. Each strain has its own intrinsic *s*_*i*_ which can give some invading strains a (typically small) systematic advantage over some or all extant strains. The inclusion of general fitness differences breaks the common assumption of ecological tradeoffs e.g. [10, 48, 49]: so if close-to-rigid tradeoffs do occur, they must emerge from the evolution rather than being assumed. Under what conditions do mutations with general (as well as specific) effects allow for or inhibit diversification?

We assume the general fitnesses are drawn from an underlying probability density, 𝒫(*s*). If a new strain, *M*, arises by mutation from a parent strain, *P*, its *s*_*M*_ is drawn from a distribution 𝒫 (*s*_*M*_*s*_*P*_) conditional on the parent’s *s*_*P*_: we choose a form for this conditional distribution that preserves the important features of the underlying distribution 𝒫(*s*). It is important that along with changes in the generalist fitness of a strain, mutations also change the interactions of the strain relative to those of its parent. (If a mutation only changed the general fitness, then the mutant would have a systematic advantage over its parent driving the latter extinct and making the evolution singular.) As a community evolves, the distribution of *s*’s of the community will change so the effectiveness of a strain’s larger *s* can diminish since what matters is its magnitude relative to the population-weighted mean

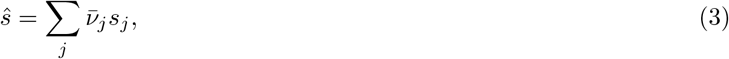

which changes under evolution.

We study a family of distributions over positive *s* of the form 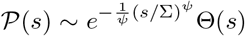 where Θ is a Heaviside function that enforces the constraint *s* > 0; Σ is a characteristic scale of the *s*’s and the parameter *ψ* controls the shape of the high-*s* tail of the distribution. [We consider only positive *s*’s but all *s*’s can be shifted by a constant without affecting the dynamics because this constant gets absorbed into the Lagrange multiplier term ϒ(*t*) to keep the population fixed.] Anomalously small *s* strains are very unlikely to successfully invade, so the sharp cutoff at the lower end does not matter.

In Figure 4 we show the results of evolutions by successive invasions of unrelated strains. For all *ψ, ŝ* systematically increases which means that only rarer and rarer strains have a chance of successively invading. The number of successful invasions, *Z*, thus only increases logarithmically in the number of invasion attempts *Y*, according to *Z* ~ log(*Y*)^*θ*^ with the exponent *θ* derived in terms of 𝒫(*s*) in Section 4.5. The evolution of the community size is seen to depend crucially on *ψ*. If the tail of the *s* distribution falls off faster than a simple exponential, *ψ* > 1, the community continually diversifies. If the *s* distribution decays slower than exponential, *ψ* < 1, the diversity decreases (after an initial increase if Σ is sufficiently small) and eventually crashes. In the marginal case of a simple exponential tail, *ψ* = 1, the diversity saturates and fluctuates around a steady state value while the mean *ŝ* increases linearly with the number of successful invasions. An analysis of these behaviors is in Section 4.4.

**Figure 4:**
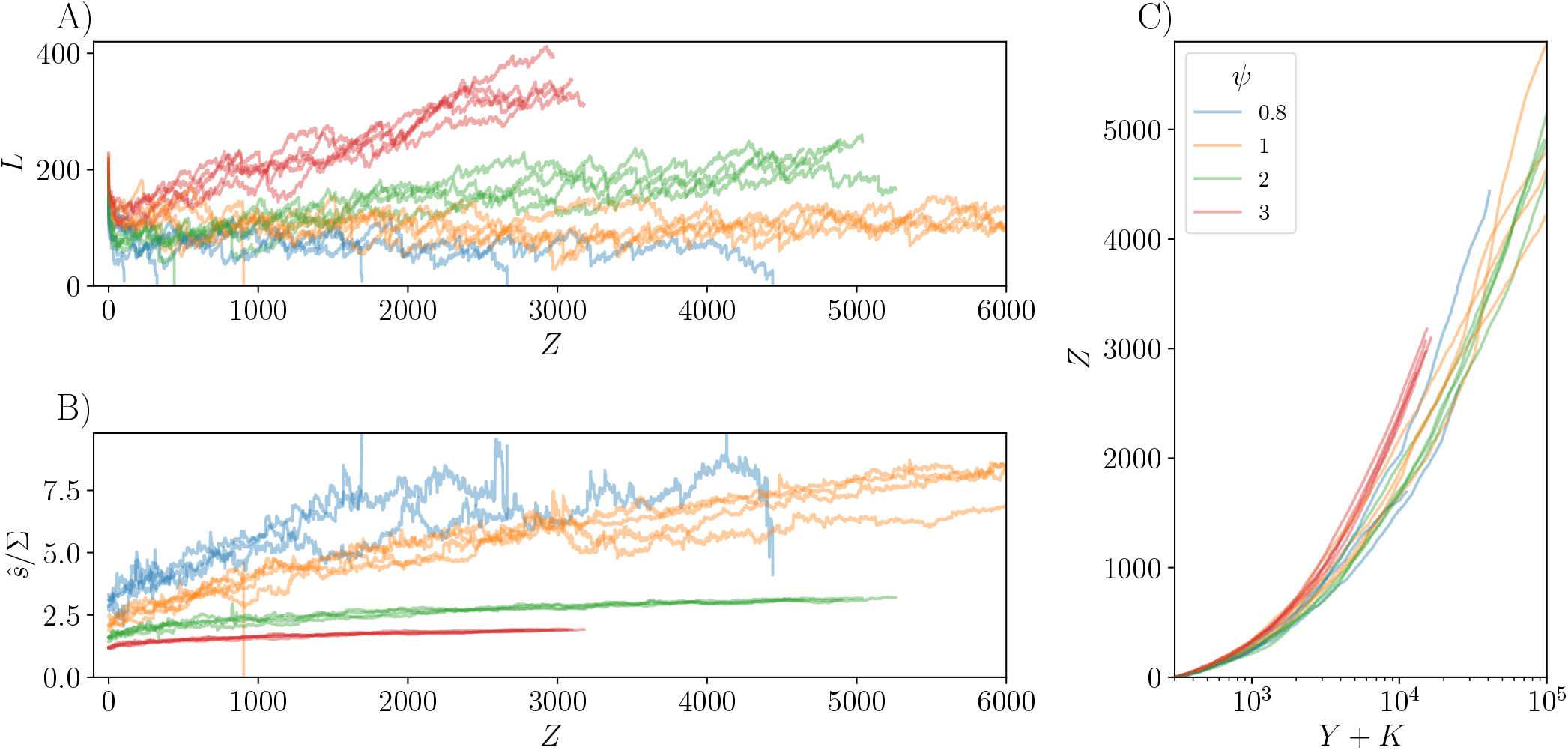
Varying the distribution of general fitnesses changes the dynamics of evolution by unrelated invaders (*ρ* = 0). (A) Larger *ψ* induces faster diversification since the distribution of invaders’ *s* falls off more steeply, meaning that differences between growth rates are smaller, driving fewer types extinct. (B) Since 𝒫(*s*) decreases faster for larger *ψ*, the community average *ŝ* increases more slowly as it becomes harder to sample into the tail of the *s* distribution as evolution progresses. (C) The rate of successful invasions *Z* versus attempted invasions *Y* shows that log *Y* is the appropriate scale for determining *Z*, since sampling deeply enough into 𝒫(*s*) to invade becomes very rare as *ŝ* increases. We offset *Y* by *K*, which is roughly the number of attempted invasions needed to buildup an initial size *K*, if nucleating into the diversifying phase were not difficult. Here Σ ≅ 0.066, 0.080, 0.177, 0.173 for *ψ* = 0.8, 1, 2, 3 respectively and *K* = 300 for each. If all the Σ were the same, the trajectories for *ψ* = 3 would diversify even faster than the others — we thus increase Σ for larger *ψ* to lessen this effect.

For *ψ* > 1, the shape of the 𝒫(*s*) is a caricature of diminishing returns epistasis. With 𝒫(*s*) decaying very sharply it is increasingly difficult for strains to produce fitter mutants as they have themselves already become very fit. Our results suggest that strong diminishing returns epistasis is conducive toward diversification, since it prevents well adapted communities from producing mutants with large general advantages that could cause the diversity to crash.

### 3.5 Exponential tail of 𝒫(*s*): *ψ* = 1

With the community-average *ŝ* growing steadily, strains with *s* substantially less than *ŝ* are very unlikely to persist, and strains with substantially larger *s* are unlikely to have yet occurred. Thus the range of *extant s*, Σ, is likely to be much narrower than Σ. This implies that the distribution over the currently-relevant range can be approximated by an exponential distribution 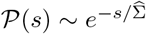 with

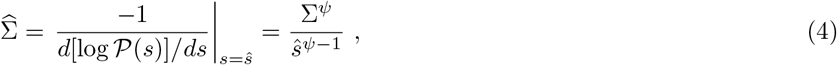

where the second equality is for the specific models we study. For *ψ* > 1, provided evolution has proceeded long enough that no strains with *s* smaller than Σ survive, 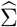 varies slowly for a range of evolutionary time and sets the scale for variations of *s*’s of extant strains and potentially-successful invaders. This suggests that understanding the general behavior can be built on understanding the case of the simple exponential distribution (for which 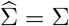).

For the simple exponential distribution, 𝒫(*s*) ~ *e*^−*s*/Σ^Θ(*s*), Σ plays a controlling role. If one strain has a substantially higher growth rate than all other strains, it will outcompete them, driving many extinct. Thus a broad distribution of magnitudes of *s* is likely inconsistent with a diverse community. We therefore focus on narrow distributions: i.e. small Σ. The typical magnitude of the bias that strains exert on each other via their interactions (the *drive* on the strain) is of order 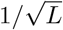. Thus the simplest expectation is that when Σ is much smaller than this, it will not matter much. On the other hand, if Σ were to be much larger than 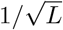, the differences in the *s*’s would dominate over the drives and only the strains with the highest and quite similar *s*’s would survive. Thus *L* ≫ 1/Σ^2^ seems inconsistent. Even for the initial community with *L*_0_ strains we expect that *L*_0_ can not be larger than order 1/Σ^2^ (although it can be much smaller if *K* ≪m 1/Σ^2^). Indeed, in Section 6.6 we show that in the limit where the community is initialized with many strains, Σ sets the initial persistent community size, *L*_0_. In the assembled community with large Σ, few strains survive the initial drop so the ones with higher fitness can reach higher abundance — which is a reason for *ŝ*/Σ increasing with Σ. This behavior can be checked in the ASM (Section 6.6).

A natural conjecture is that for small Σ with an exponential distribution, steady diversification can occur until the breadth of the *s* distribution becomes important — when *L* ~ 1/Σ^2^ — and after that *L* will saturate, as seen in Figure 5. Thereafter *ŝ* will grow and invasions of unrelated strains are less and less likely to be successful; but when they are, on average they will drive exactly one other strain extinct. Figure 5 illustrates this behavior, including the large initial drop from *K* to *L*_0_ when *K* ≫ 1/Σ^2^.

**Figure 5:**
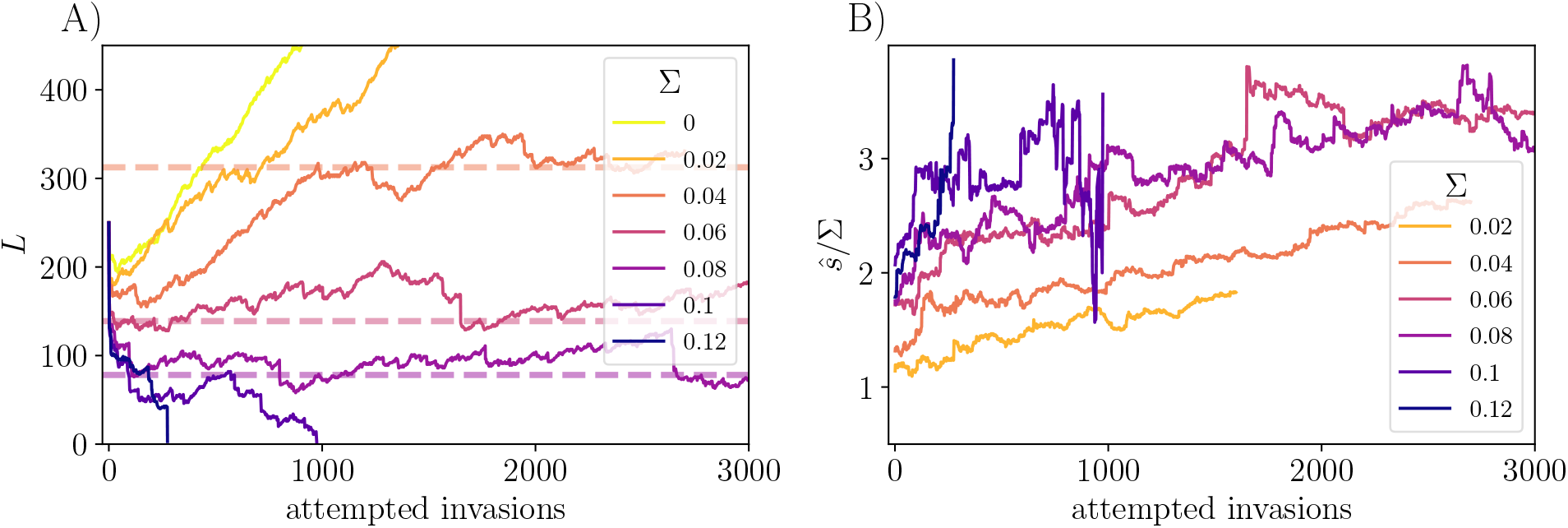
Effect of exponentially-distributed *s* on community evolution. (A) When 𝒫(*s*) ~ *e*^−*s*/Σ^ (*ψ* = 1), there is an evolutionary steady state with *L* ~ Σ^−2^. For a narrow distribution of the general fitnesses (*L* ≪ Σ^−2^), *L* increases linearly before saturating. For larger Σ with many initial strains, immediate extinctions drive *L* down to *L*_0_ ~ 1/Σ^2^ (Section 6.6). The dotted lines indicate Σ^−2^/2 for the intermediate values of Σ for which saturation is observed. (B) The average fitness of the community, 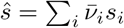 (shown scaled by Σ) grows linearly with the number of successful invasions — but the rate of successful invasions decreases. For the simulations that reach steady state *L*, the rate of fitness increase per successful invasion, *dŝ*/*dZ*, scales as Σ^3^ (Figure 15).

Armed with this understanding of the exponential case, *ψ* = 1, we can understand the behavior of the super-exponential case, *ψ* > 1. When the initial *L*_0_ ≪ 1/Σ^2^, the diversity will increase linearly in evolutionary time, *Y*, until *L* ~ 1/Σ^2^ and *ŝ* ~ Σ. Beyond this point the evolution will gradually slow down with *ŝ* gradually increasing. The invaders that have a substantial chance of success will have a distribution close to a simple exponential with scale, 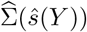, given by Equation 4. The community size will be 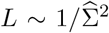 with the same coefficient as the simple exponential model, and the joint distribution of *s* − *ŝ* and 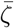 of the extant strains will become closer and closer to that of the simple exponential model. We analyze this behavior and find the diversity trajectory *L*(*Y*) in Section 4.5.

For a stretched-exponential tail of 𝒫(*s*), *ψ* < 1, as *ŝ* increases 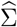, the width of the distribution of extant *s*_*i*_, will broaden and concomitantly we expect that the diversity will decrease. Eventually — in practice rather soon — *L* will decrease enough that it will crash. However if 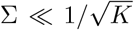, the diversity will initially increase and reach a maximum before decreasing and crashing as the effects of the tail of the distribution of *s* are felt. We conclude that for the evolutionary process in these models to *continually* generate higher diversity, the distribution of general fitnesses must decay sufficiently rapidly. (Note that with a rigid upper bound to *s*_*j*_, the diversity would continue to increase while slowing down less dramatically than the models with no upper bound.)

### 3.6 Correlated general fitnesses

We now briefly illustrate some of the combined effects of mutations and nonspecific advantages. These are complicated in ways we only partially understand, and there are several crossovers that make interpreting the simulations difficult. For the conditional distribution of mutant *s*_*M*_ given parent *s*_*P*_, we make a simple gaussian approximation which preserves the form of the important high *s* tail. We take 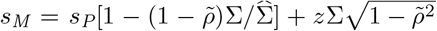 with *z* a standard normal random variable, but truncated so that all *s*’s are positive, and 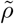 is a measure of the degree of correlation between *s*_*M*_ and *s*_*P*_. In the case of *ψ* = 2, on which we focus for now, this defines the conditional distribution

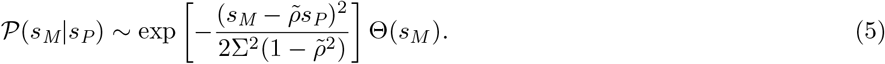

(Although the marginal distribution of *s*_*M*_ is not exactly a truncated gaussian, as 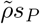 becomes larger than 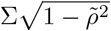, the effects of the truncation will not be significant, and the tail of 𝒫(*s*_*M*_) remains gaussian, as desired.) In general the degree of correlations, 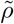, between the general fitnesses, and the correlations between the mutant’s and parent’s interactions, parametrized by *ρ*, can be different. But for concreteness we take 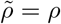.

Figure 6 shows the results of evolutions with *ψ* = 2 over a range of *ρ*. Correlations increase the rate of growth of *ŝ* since the descendants of strains with high *s* have a higher chance of having even higher *s* because of the correlations. However if *ρ* is very close to 1 then the rate of increase of *ŝ* slows down simply because the increments in *s* of the mutants become too small to increase the mean substantially. Therefore the rate of increase of the population-mean *ŝ* is highest for an intermediate value of *ρ*.

**Figure 6:**
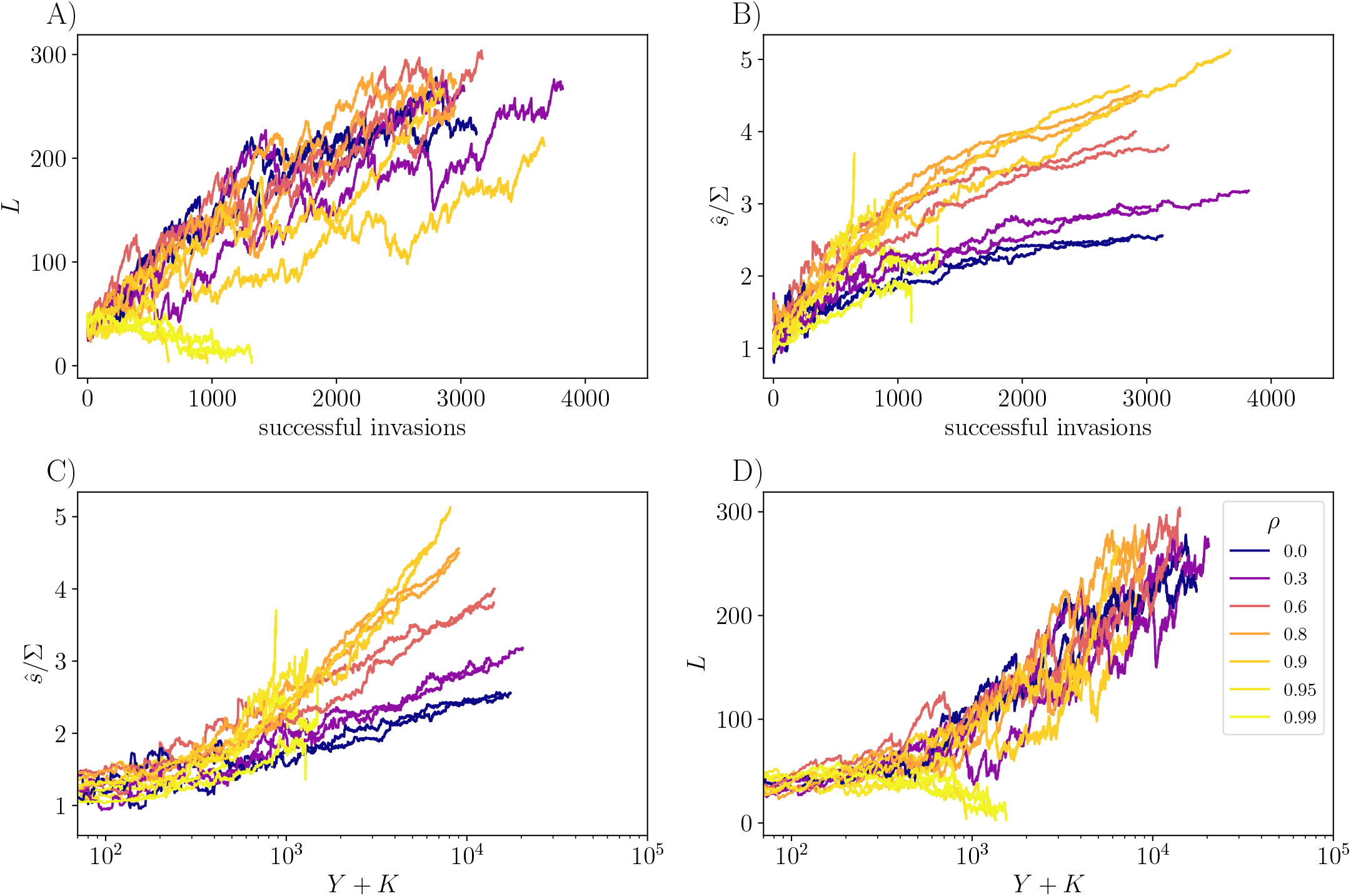
Correlations between parents and invading mutants modify the effects of general, nonspecific differences. Here, the distribution of *s* of the *K* = 50 initial strains is a half gaussian (*ψ* = 2 with Σ ≅ 0.177) (A) Trajectories are shown conditional on not crashing, except for *ρ* close to unity for which crashing from *K* = 50 is exceedingly likely. There are competing effects caused by the correlations. Larger *ρ* means it is easier to invade because the mutant *s*_*M*_ is closer to the parental *s*_*P*_, but correlations make it more difficult to coexist because the mutant’s interactions are very close to those of its parent. The net effect is that curves of *L* versus *Y* are quite similar for different *ρ*. (B) Correlations make *ŝ* increase faster, since the average increase in *s* conditioned on invasion is higher. The ratio of successful to attempted invasions is higher with mutant-parent correlations, as is more evident in Figure 11. (C) and (D) same data as (A) and (B), but versus attempted invasions, *Y* (for which the *x* axis on a log scale is more natural).

Though we have confined exploration of correlated general fitnesses to the case of *ψ* = 2, we expect similar behavior for all *ψ* > 1. By contrast, for *ψ* < 1, the community will crash as in the absence of correlations.

## 4 Analysis

In this section we develop an approximate analytic theory of the evolutionary dynamics and provide heuristic understanding for most of the observed phenomena described above. The underlying basis is the dynamical mean field theory (DMFT) of the STC phase developed in PAF. This takes advantage of the large number of strains to simplify the descriptions and analyses of the behaviors.

The natural quantities that characterize strains in the DMFT are their biases, {*ξ*_*i*_}, and how these set their mean abundances, 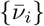. For a large randomly assembled or evolved community the mean abundances will be a function of the biases,

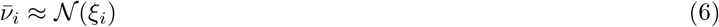

with 𝒩 depending on the parameters and evolutionary history. The relation between 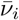 and *ξ*_*i*_ and the *drive* the other strains exert on *i*, enable one to estimate the bias from the simulations (Section 6.7). Armed with the DMFT description, we can understand how the biases of extant strains change over the course of invasions. We do this in detail for the simplest case of invasions of unrelated strains without general fitness differences and show that the evolution causes *L* to change linearly with the number of invasions — decreasing or increasing depending on the parameters. A simple approximation to the evolution enables semi-quantitative results, including for the scaled distribution of biases that is achieved for *L* large and steadily increasing.

We then analyze the effects of general fitness differences first with a simple exponential distribution of *s*, 𝒫(*s*) ∼ *e*^−*s*/Σ^, and show how the scale Σ gives rise to a saturating value of *L* as in Figure 5. Based on this, we generalize to other shapes of the high-*s* tail, parametrized by *ψ*, showing how the steepness of the tail of 𝒫(*s*) affects the rate at which *L* increases or decreases.

Building on the understanding of invasions of unrelated strains, we then analyze, mostly heuristically, the effects of correlations, of magnitude *ρ*, between mutants and parents, studying both interaction and nonspecific mutations.

Finally we compare the properties of ecologically and evolutionarily stable communities, and find that conditioning on gradual emergence of diversity leaves only subtle and small signatures on the measurable properties of the community.

### 4.1 Dynamical mean field theory

The DMFT approximation, which is exact in the limit of a large number of strains with random interactions between them, replaces the full statistical dynamics by the stochastic effects of the others on one chosen strain, with the statistical properties then determined self-consistently from the properties of the distributions over the strains. This approach was first used in the physics of disordered systems such as spin glasses [50], but has been applied to ecological dynamics in a number of subsequent works [42, 43, 44, 26, 51, 40].

When a strain, say *i*, has very low abundance, the effects of it on the others are very small so the forces of the others on it, Σ_*j*_ *V* _*ij*_*ν*_*j*_, are a sum of roughly independent random variables and thus act like gaussian noise with correlations *C*(*t, t*′) = ⟨Σ_*j*_*ν*_*j*_(*t*)*ν*_*j*_(*t*′)⟩. But when it rises to substantial abundance, it will weakly affect the others: e.g. *ν*_*j*_(*t*) will change by an amount 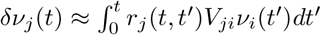 where *r*_*j*_ (*t, t*′) is the linear response of *ν*_*j*_ (*t*) to a change in its growth rate at an earlier time *t*′. The cumulative effects of these changes feed back onto strain *i* to modify its growth rate at a later time *t* by Σ*V*_*ij*_*δν*_*j*_ (*t*). Because of the correlation of strength *γ* between *V*_*ij*_ and *V*_*ji*_, the terms add coherently resulting in a modified growth rate of *i* by 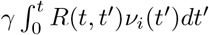, with *R*, determined by feedback from the total impact on the community of the strain’s own past history. This results in autonomous stochastic integro-differential equations for each strain independently: as they are statistically equivalent on each island, we drop the island subscript:

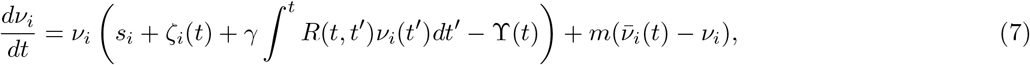

with the gaussian *drive ζ*(*t*) with correlations *C*(*t, t*′), the memory response *R*(*t, t*′) = Σ_*j*_ *r*_*j*_(*t, t*′), and the island-averaged abundance 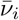, to be determined self-consistently as in PAF. Note that we have assumed here that the correlation between *V*_*ij*_ and *V*_*ji*_ and their variances do not change substantially in evolved communities: we show in Section 4.8 that our simulation results are consistent with this *Ansatz*.

The growth rate of strain’s population on an isolated island is the sum of its general fitness *s*_*i*_, the time-dependent drive *ζ*_*i*_(*t*), the feedback part, and the time-dependent Lagrange multiplier, ϒ(*t*). In ecological steady state, some of the strains will have gone extinct, and the correlations and responses of the persistent strains will only depend on time differences *t* − *t*′, and quantities will have well-behaved time averages equal to island-averages, denoted by overbars. From Equation 7 we see that the average population growth rate of strain *i* when it is at *low* abundance, its bias, is *ξ*_*i*_ = *s*_*i*_ + ⟨*ζ*_*i*_⟩− ⟨ϒ⟩. (This is equivalent to the prior definition in terms of the community in the absence of *i*, but now in terms of the emergent mean field quantities.) The average growth rate of strain *i* on a single island excluding migration is

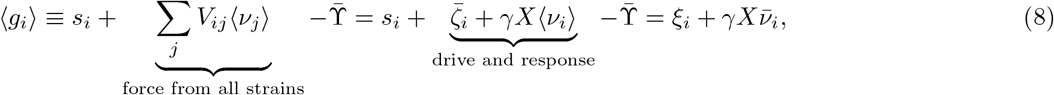

where the (static) susceptibility, *X*, is the time-integrated total response function 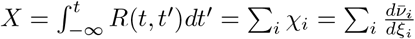 with the individual static susceptibility of each strain *χ*_*i*_ = ∫*r*_*i*_(*t*)*dt*. In steady state, 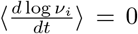 so that ⟨*g*_*i*_⟩ < 0 must be exactly compensated by the average effect of the migration term 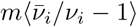, which is always positive (by the Cauchy-Schwarz inequality) and can be much larger than *m* when *ν*_*i*_ is small.

The stochastic DMFT equations cannot be reduced to equations for the correlation and response function and must be treated directly. Their self-consistent solutions in the STC phase were analyzed in our previous work [PAF], by asymptotics in the large parameter *M* = log(1/*m*). As discussed in PAF this yields super-diffusive random walks of the log-abundances, persistence around the migration floor, and the statistics of the occasional blooms that occur for all persistent strains with negative bias. These aspects together determine how the mean abundance of a strain depends on its bias, via 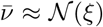 with corrections smaller by factors of 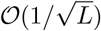.

When the bias of a strain is positive its mean abundance will be quite large, the average input from migration relatively small, and hence ⟨*g*_*i*_⟩ ≈ 0 which implies that ⟨*ν*_*i*_⟩ ≈ *ξ*_*i*_/(−*γX*). For strains with negative bias, the migration is essential and the statistics of the blooms, which are rarer the more negative the bias, make 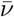 exponentially small in −*ξ* [PAF]. For strains with large negative bias, *ξ* < *ξ*_*c*_ < 0, the migration cannot sustain their local populations and they go globally extinct with 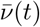 decaying exponentially in time (in the deterministic approximation which of course breaks down as they go extinct.)

The distribution of the biases of a community plays a crucial role. In an initially randomly assembled community of unrelated strains, the biases are essentially independent (up to corrections smaller by 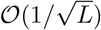) with the mean drives, 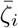, being gaussian distributed with average zero and standard deviation of order 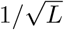. More precisely, since *ξ*_*i*_ is determined by the community in the absence of *i*, the *V*_*ij*_ that determine it are independent of that community. Hence even if extinct strains are included, the variance of 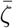, and thereby also of *ξ*, is simply 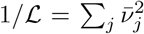 (with only small corrections from the neglect of the strain itself): this simple result is explicit in the self-consistency condition of the DMFT correlation function. We therefore define ℒ as an *effective size* of a community which weights the contributions of strains by their mean abundances and, unlike *L*, is insensitive to whether the close to marginal strains have or have not gone extinct. The distribution of the biases of the persistent strains in the randomly assembled community is therefore a truncated Gaussian with lower limit *ξ*_*c*_, which is of order 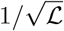 — the magnitude of *ξ*_*c*_ and ℒ must of course be determined self-consistently. Crucially, these will depend on the dynamics — especially the blooms — not just the mean quantities.

With the dynamical mean field understanding of an assembled STC phase in hand, we can proceed to describe the evolutionary process in terms of the distributions of properties of the extant and newly invading strains — of particular importance their biases and consequent mean abundances. The distribution of biases is perturbed by the introduction of new strains and this can push some of the extant biases below *ξ*_*c*_, which itself depends on the bias distribution as modified by prior evolution, and on the number of extant strains, *L*. Importantly, once a strain goes globally extinct, it cannot emerge again, even if the conditions become favorable for its reappearance: Without permanent extinctions, the evolutionary process would be just a gradual assembly of strains and would arrive at the same final state that would occur if all the strains that emerged over the course of the evolution were placed into the environment together.

Our analysis and description here is in the limit of infinite number of islands, but the simulations are done with *I* = 40 islands, which requires us to justify drawing conclusions from this modest number of islands. For an assembled STC phase, we showed how rare fluctuations to extinction occur for large but finite *I* with rate exponentially small in the number of islands. For a strain with bias, *ξ*, the extinction rate is of order exp[−*I*/*I*_*X*_ (*ξ*)] with the characteristic island number, *I*_*X*_, depending on the bias. For close to marginal strains, *I*_*X*_ is quite large so extinctions will happen that would not have with *I* = ∞. But these marginal strains would, in any case, have little effect on the extant strains: Thus their loss should not matter much — particularly true if we use the effective size ℒ, which gives little weight to low mean-abundance strains, instead of *L*. For strains further from marginal, the time the simulations are run for in each epoch (between successive invasions) is modest enough that these strains are unlikely to fluctuate to extinction.

### 4.2 Evolution without general fitness differences

We first analyze invasion of unrelated strains without general fitness differences. As the bias of an attempted invader *A*, given by 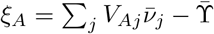, is determined by its interactions, {*V*_*Aj*_} with the extant strains, this drive part is gaussian with mean zero and standard deviation of 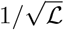 *independent* of correlations among the extant strains (although these will affect the 𝒩(*ξ*) and hence ℒ).

A new strain *A* can successfully invade the community and persist if it has, *ξ*_*A*_ > *ξ*_*c*_. The probability of successful invasion is thus simply 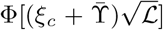, with Φ the standard normal cumulative distribution function. In the initially assembled community, 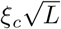 and 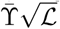 are independent of ℒ. If we make the *Ansatz* that after a long period of evolution the scaled statistical properties of extant strains reach a steady state — albeit a different state than the initially assembled community — then the invasion-success probability will become independent of *L* for large *L*. If the mean number of extinctions per successful invasion also reaches a steady state value, which is less than unity, this explains the steady linear growth of *L*(*Y*) with attempted invasions seen in Figure 2.

However we need to understand the effects of invaders on the extant strains, and explain the observation, in Figure 3, that for *L* ≫ *L*^*^, the number of extinctions per invasion is roughly exponentially distributed with mean less than 1. This requires understanding the evolutionary dynamics of biases under successive invasions. With the one-by-one introduction of new strains, the biases of each extant strain undergoes some kind of random walk, and the strain goes extinct if its bias ventures below *ξ*_*c*_. In Figure 7A we show the evolutionary trajectories of the biases of 5 individual strains that started from similar initial values in a simulation where the community diversified from 50 to 500 strains. Extinctions are caused by *ξ*_*i*_ being pushed to below *ξ*_*c*_ by an invading strain. (Note that for finite *L*, the effective sharpness of *ξ*_*c*_ will be smeared by and amount of order 1/*L* due to variability in the dynamic noise from strain to strain which we have not explored. Only for *L* very large, is there is a sharp boundary at the scaled bias 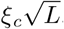.)

**Figure 7:**
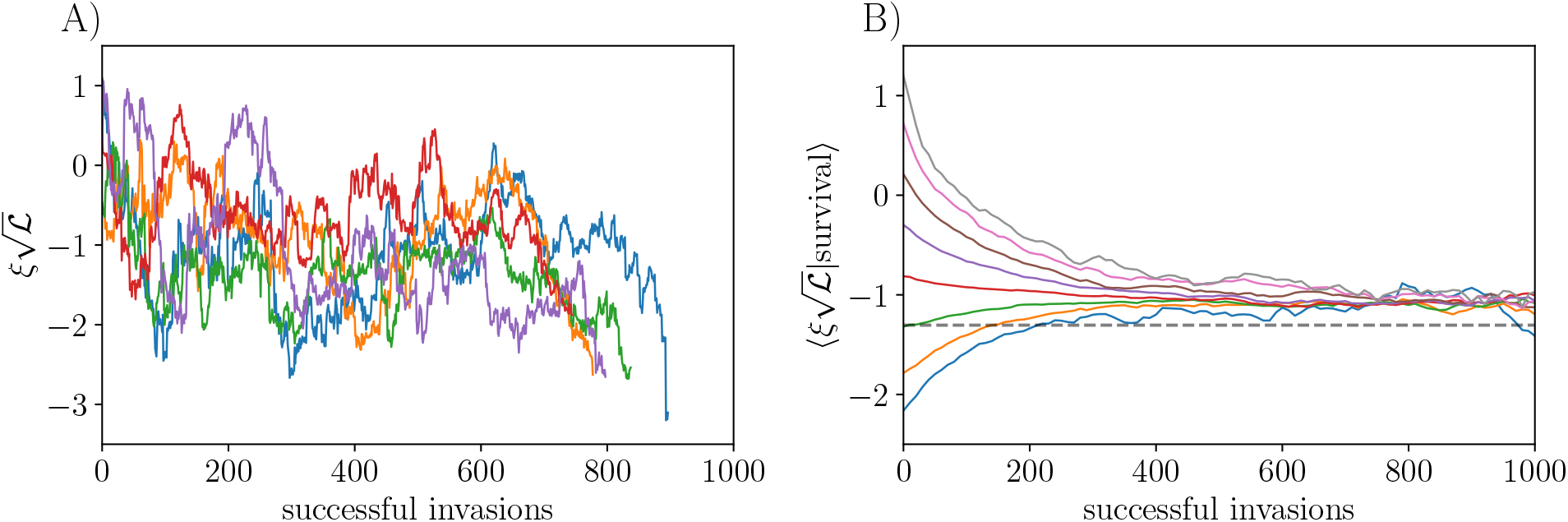
Trajectories of biases of persistent strains under the influence of successive unrelated invaders with all *s*_*i*_ = 0. (A) Bias trajectories (normalized by 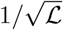, the standard deviation of the invader’s bias) of individual strains that persisted in the community for between 800 and 1000 subsequent successful invasions after themselves invading at different times during the evolution. Extinctions occur when a trajectory terminates, and result from the bias going below the critical bias. (B) Bias trajectories for all strains binned into groups by their starting value and averaged within bins for as long as the strains persist. Without conditioning on success, the biases of new invaders have mean 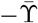 and standard deviation 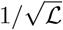. The dashed line shows 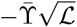. Conditioned on survival, the (normalized) biases converge to and fluctuate around a larger value. Data in (A) are from a single simulation where the community diversifies from 50 to 500 strains, and in (B) data are pooled from 10 replicates of the same process.

To further investigate the nature of the bias evolutions, we average over a large number of strains, binning them according to their initial values normalized by 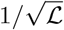. We observe a strong tendency of anomalously positive and negative biases to regress toward an intermediate value. In this plot, as evolution proceeds, the asymptotic average bias conditioned on survival is larger than 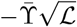. (Figure 7B). This is because of the conditioning on survival of the strains: those that persist for many epochs tend to have larger-than-average (but still negative) bias.

The stochastic evolutionary dynamics of the biases is subtle because the current community is conditioned on, and correlated with, its full evolutionary history. In principle one could carry out a “dynamic cavity” method analogous to that used to derive the DMFT for an assembled community [26]. To do this, one would follow the bias history of a *probe strain*, labeled 0, which never actually entered the community and thus does not affect it. The interactions {*V*_0*j*_} affecting the probes are thus independent of the community. From the statistics of the bias trajectory of the probe, *ξ*_0_(*Y*), and knowledge of the relationship between the bias and mean abundance at each 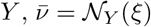, one could infer the trajectory of the abundance the probe strain would have had from when it first successfully invaded — if it had — until when it would have gone extinct when *ξ*_0_(*Y*) dropped below *ξ*_*c*_(*Y*) for the first time. With the ensemble of the trajectories of the mean abundances of invaders from all earlier times through their persistence and extinctions thus constructed, one could use the basic property of DMFT (in the large *L* limit), that the dynamics of all the strains are statistically equivalent to those of a probe strain. As the statistics of the bias of the probe strain across evolutionary time, *Y*, are gaussian with mean 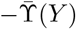 and covariance between two evolutionary times given directly by the correlation function 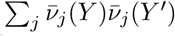, the self-consistency condition on the full evolutionary statistics could in principle be used to “solve” for the needed quantities and distributions. Of course one would need to also know the function 𝒩_*Y*_ (*ξ*) which could be obtained numerically once one knows the distribution of biases for each *Y* : together these would need to be determined self-consistently. We have not attempted to carry out this program (which is analogous to that done to analyze the spatiotemporally chaotic dynamics of the assembled community in PAF). Instead, we analyze a simple approximation to the random walk of a probe strain and the correlations induced by the evolution. For this, there are only a small number of unknown constants which could more readily be obtained from simulations.

We make a Markov approximation of the effects of the evolutionary history and assume that the distribution of the biases at epoch *Y* + 1, only depends on their own values at epoch *Y* and the overall properties of the community. Consider the drive (here meaning the “mean-drive”) of strain 0 before, 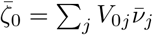, and after,

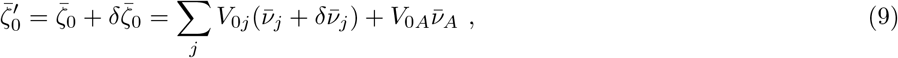

the successful addition of a new strain *A* to the ecosystem which establishes with mean abundance 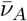. The sums on *j* do not include 0 or *A* and the δ’s denote changes that result from the invasion. As 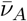 is of order 1/*L*, its *direct* effect on the bias is smaller by a factor of 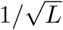 than 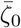. However *A* will also change all of the other biases by similar-magnitude random amounts with uncorrelated signs, causing 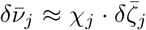. Thus 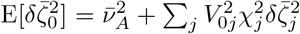 with the expectation over the interactions with the probe strain. Since the properties of the extant strains depend neither on the *V*_0*j*_ nor the perturbations via *V*_*jA*_, the averages over each of the squared factors can can be performed separately leaving 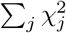 times the average over *j* of 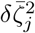. But now we can consider each strain separately to be the probe strain so that the average over the extant strains must be the same on both sides. This yields 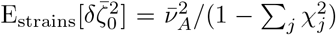. The more directly measurable quantity is the total squared abundance change of the extant strains caused by a random change in growth rate of each: With the perturbation caused by the invader, we have 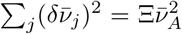 in terms of the nonlinear response — which we call the 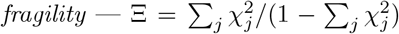 (This is analogous to the spin-glass susceptibility of random magnets [52]). The fragility is of order unity and the form of the denominator shows that it can diverge: such divergence indicates an instability of the community (and breakdown of the DMFT *Ansatz*). The fragility weights more heavily the higher abundance strains as the change in the absolute abundance of the close-to-marginal strains is small even if their abundance changes by a substantial multiplicative factor.

[Note that the fragility is a general measure of the sensitivity of a community to perturbations. In the stable niche-phase for large *Q*, the fragility diverges as *K* increases to the stability boundary at *K*_crit_ ~ *Q*^2^. It is infinite in the ASM which is exactly marginal and highly sensitive to added strains.]

In addition to the random change of a 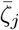 induced by a successful invader, there is also a systematic change, as seen in Figure 7. This is because 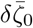 and 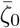 involve the same set of random *V*_0*j*_ and are thus correlated. As they are both sums of independent gaussian variables times coefficients, averaged over the *V*_0*j*_ the conditional expectation is 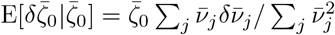. But since each 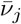 and 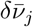 are correlated as a consequence of 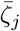 and 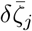 being correlated, we again have to self-consistently determine this correlation. Unfortunately this is more complicated as the 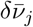 also have a contribution from small changes in the function 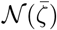 caused by the invaded strain. However one can understand the *form* of the conditional expectations by noting that after the new strain has invaded, 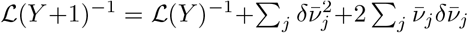. This implies that the last term must be negative if *L* and ℒ are increasing or staying constant. Note that the correlation between 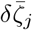 and the current 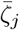 caused directly by the interaction of the invader on strain *j* is, indeed, generally negative. The additional effects on *j* from the changes in other strains is enhanced by the fragility, Ξ. If the fragility is high, one can show that this conditional expectation is proportional to −Ξ: in this limit, as argued below, the diversity will decrease.

The above Markovian analysis is, as already said, not correct because really 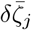 for each extant strain is correlated with its whole past trajectory — as it involves many of the same *V*_0*j*_ — and its conditional expectation will be a weighted sum over the full past 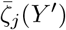. Furthermore, it is possible that conditioning on all this past could suppress 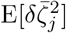 by a substantial factor (in the Markovian approximation the analogous correction is smaller by a 1/*L* factor). However the basic structure of the stochastic changes is correctly captured by the Markovian approximation.

Both the systematic and the random change in the drive are proportional to the average abundance of the invading strain, which is of order 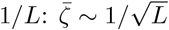 and 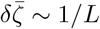 while its conditional mean is of order 1/*L*^3/2^. These scalings mean that in the Markovian approximation there is a natural phenomenological Langevin equation for the changes of the drives due to invasions:

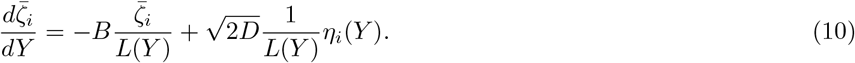

with *η*_*i*_(*Y*) approximately gaussian with mean zero and unit variance — an approximation that should be good if we average over a substantial range of *Y* (but with range much smaller than *L*). From the above analysis, we saw that *B* and *D* are order-unity coefficients which respectively characterize the average and mean-squared response of the bias to the invasion of a new strain. They are both proportional to 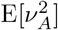 times the fragility, Ξ, of the extant community. This Langevin equation must be supplemented by a boundary condition that if 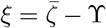 goes below *ξ*_*c*_, the strain disappears. If neither extinctions nor invasions occurred, there would be a trivial steady state with a gaussian distribution of the biases with width 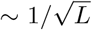 as expected for an assembled community. With a source of new strains, which come in with drives that are gaussian distributed with width 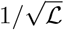, the effective community size is ℒ ≈ *CL* with *C* < 1. We thence obtain a Fokker Planck equation for the number density *N* of the drives:

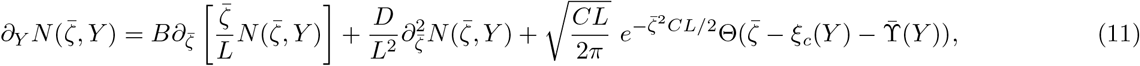

where we impose the absorbing boundary condition at the critical bias corresponding to 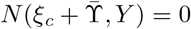 with *ξ*_*c*_ and 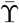 both scaling as 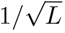. The last term in Equation 11 is the truncated-gaussian drive distribution of successfully invading strains with scale 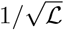. The fraction of invasions that are successful, *dZ*/*dY*, is just the integral of the truncated gaussian, 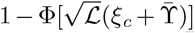. The number of strains, *L*(*Y*), is changing and at this point unknown: it is determined self-consistently from 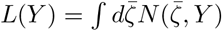. It is straightforward to show, as we do in Section 6.8, that this Fokker-Planck equation admits a scaling solution with the *Ansatz* that all the quantities scale as 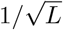. The rate of diversification, *U* = *dL*/*dY*, is of order one and determined by an eigenvalue-like condition: it can be either positive or negative depending on 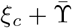 and the coefficients, *B, D, C*. If the community is very fragile (Ξ large), then the *B* and *D* terms will dominate over the input and the loss of strains per successful invasion will be large: in this regime the diversification rate is negative. On the other hand if 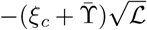 is large, few strains will go extinct and the community will diversify. Of course, these quantities are determined self-consistently, depending on both the distribution of biases and the function 𝒩(*ξ*), which itself will evolve, reaching a scaling form in the steadily diversifying state.

### 4.3 Evolution of the bias distribution and distribution of number of extinctions

The approximate model of the evolution of the biases makes predictions about the shape of the bias distribution as a function of time. Before the onset of evolution, the distribution of biases is a truncated gaussian, with a low end cutoff set by *ξ*_*c*_. However as the distribution evolves, approximately under Equation 11 with the absorbing boundary condition at the critical bias, it smooths out near this cutoff, going linearly to 0 as 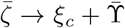 (or, equivalently, as *ξ ξ*_*c*_). This linear density of states at the low end of the bias distribution determines the response of the community to evolutionary perturbations, since these low-bias strains are the ones most susceptible to extinction. In particular, this linear density of biases determines the distribution of the number of extinctions per successful invasion (Figure 3B).

To estimate the distribution of the number, 𝓁, of strains that goes extinct when there is an invasion — particular the probability that 𝓁 is large — we note that the effect of an invader will be to perturb the extant strains’ biases by some random amount of order 1/*L* — thus much smaller than the width of the bias distribution — and proportional to the mean abundance of the invader. (The average 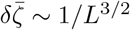 is much smaller than the random part, as already noted.) The positive tail of the invader’s mean abundance, 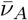, is gaussian, since for positive 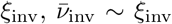 and the invader’s bias *ξ*_inv_ is itself gaussian distributed. The number of strains whose biases are within 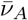 of *ξ*_*c*_ is proportional to 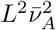, because the distribution of strains bias’s vanishes linearly at *ξ*_*c*_. Thus for fixed 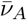, the number of strains that are driven extinct is Poisson distributed with mean proportional to 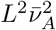: this is of order one for large *L* as expected. That the tail of the distribution of 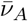 is gaussian implies 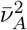 is approximately exponentially distributed in its tail. Integrating the Poisson distribution over this yields, for large 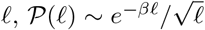 (with *β* an order-unity coefficient) which is close to exponential as observed in Figure 3B.

A similar analysis for the initial randomly-assembled community shows that for fixed 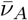, the mean number of extinctions triggered by the first successful invasion is of order 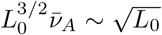; much larger than after evolution has proceeded for a while. As the Poisson with this mean has a narrow distribution, the probability of an anomalously large number of extinctions will be dominated by the gaussian tail of the distribution of 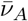 and hence itself be roughly gaussian. (However, unless the initial *L*_0_ is huge, the tail is unlikely to still be in the asymptotic regime.) The transient caused by a set of early invasions will likely cause a total of order *L*_0_ strains — with a relatively small coefficient — to go extinct before *L* starts steadily increasing, and this will occur over of order *L*_0_ invasions. For *γ* = −0.8 this effect appears to be very small —the critical bias is quite negative — but for smaller or larger *γ* the effects are noticeable (Figure 2).

Simulations exhibit the scaling relationship between the width of the bias distribution and the number of strains (Figure 8A) for evolutions over one order of magnitude in *L*. One observes a smoothing out of the bias distribution toward the low end when comparing the evolved communities that have evolved up to 500 strains from *K* = 50, to assembled communities that have *L*_0_ ≈ 500 after an initial drop from *K* = 650 (Figure 8B). However the bias distributions are very similar except near *ξ*_*c*_. This is what one expects from the Markov approximation: for 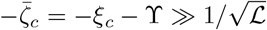, when the ratio of extant to invading widths, 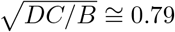, is close to unity, the bulk of the drive distribution is close to gaussian with mean zero, which is reflected in Figure 8B.

**Figure 8:**
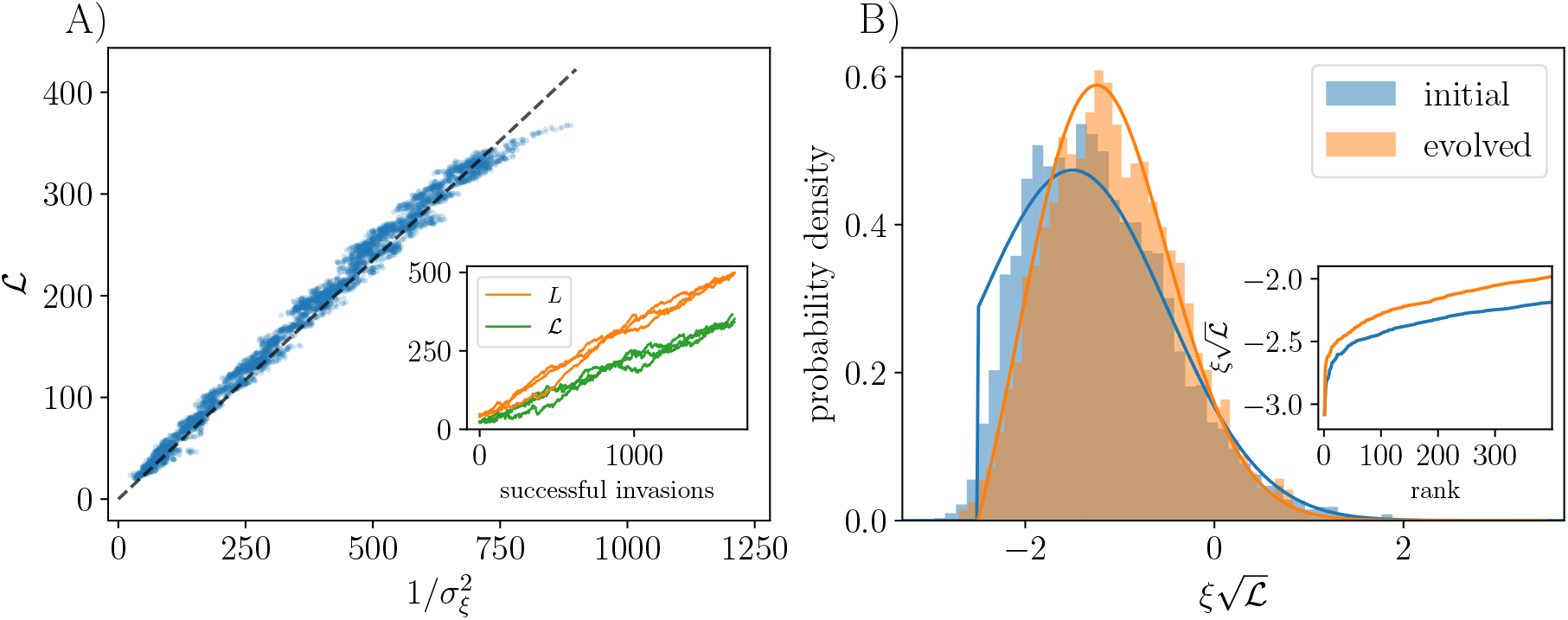
(A) Scaling relationship between the effective community size, 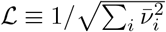, and the inverse variance of the bias distribution, 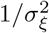, for unrelated invaders with *s* = 0. Data are pooled over 3 runs from *L* = 50 to 500 strains (inset) so that when *L* becomes large the initial conditions have been forgotten. Inset shows that *L* and ℒ are proportional as expected. (The bend in ℒ versus 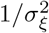 toward the end of the curve is likely due to stopping the simulation the first time *L* reaches 500, which truncates the curve asymmetrically.) (B) The low end of the bias distribution changes from a truncated gaussian for the initial involved community (blue), to a linearly vanishing function (orange) because of evolution-driven extinctions of close-to-marginal strains. Bars show histograms from simulation, and solid lines show theory. The blue curve is a truncated unit-variance gaussian for the initial assembled community, with mean and lower limit fit by hand, and the orange curve is a numerical solution to Equation 41 with *U* = *dL*/*dY* ≅ 0.24 and *B* = 2.11, D = 1.79 adjusted to enforce normalization. Note that *C* = ℒ/*L* ≅ 0.74. The mode of the truncated gaussian is 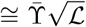 as expected from Figure 7. Although *L* is not really large enough to see the difference between truncated and a linearly-vanishing form (due to variability in the effective strain-dependent critical bias being smaller than *ξ*_*c*_ by only of order 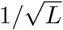, and to the low probability density in the tail) the effect is still noticeable. Inset shows the normalized bias by rank order, illustrating the smoothing of the lower end of the distribution caused by the evolution. For histograms and inset, data are pooled across 10 runs. In order to appropriately compare them the unevolved assembled communities have *K* = 650 so that *L*_0_ ≅ 500.

How good is the Markov approximation? The average change in the drive should really be a weighted integral over the history of the drive, 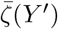, over all *Y* ′ < *Y*. If one makes the *Ansatz* of a scaling solution with *L* steadily increasing, then for large *Y, L ∝ Y* and in the variables scaled by *Y* the weighting function should be a function of of *q* = log(*Y*/*Y* ′), so that the integral over the history is a convolution in *q*. In general, the steady state could, of course, not be found even if this weighting function were known, and the integro-differential generalization of the Fokker-Planck equation would have to be analyzed numerically. But with the widths of the distributions of the extant and invader biases being similar, as observed, the details of the weighting function might not much affect the bias distribution.

### 4.4 Exponential distribution of general fitness: *ψ* = 1

As a starting point to understand the effect of nonspecific mutations on the evolution of the bias distribution, we consider exponentially distributed general fitnesses: as discussed earlier, we expect that this will serve as a basis for more general understanding of the effects of the high *s* tail of the distribution. We first analyze unrelated invaders.

A heuristic understanding of how the dynamics of *L* depend on 𝒫(*s*) follows from the mean field relation between the distribution of biases and *L*. Without general fitness differences, the biases *ξ*_*i*_ are distributed with characteristic scale 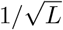. As *L* increases, the distribution of these biases gets tighter around the origin: the system becomes progressively more “neutral” with overall differences in strain growth rates becoming smaller. The contribution of the general fitness differences is to add an extra piece to each bias, broadening the distribution of extant biases. In the limit of many invasions, the width of the drive distribution becomes comparable to the width of the distribution of the *extant s*’s and cannot decrease further, as explained earlier (Section 3.5). Therefore after many invasions, the shape of 𝒫(*s*) determines both the width of the bias distribution, and the number of coexisting strains.

We can develop intuition about the effect of exponentially distributed general fitnesses by first asking how they impact the fraction of strains that survives in a randomly assembled community. In Section 6.6 we show analytically that when *K*Σ^2^ ≫ 1 with an exponential distribution, the number of persistent strains in the totally antisymmetric single-island model (ASM) is Σ^−2^ with coefficient 1. Although with multiple islands the spatiotemporal chaos will modify this relation, since it enables strains with negative bias to persist, these effects will change the coefficient but not the scaling of the number of persistent strains with Σ.

What happens if the initial number of strains *K* is smaller than Σ^−2^? Here the width of the initial bias distribution will be set by 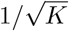 rather than Σ, and the general fitnesses will not affect the initial ecology-caused drop in the number of strains from K to L_0_. At first, *L* will increase under successive invasions. However Σ will start to matter as the distribution of drives 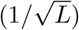 becomes narrower. The scale of 𝒫(*s*) sets a lower bound on the scale of the biases, thereby fixing a value of *L* at which the community diversity saturates: As the distribution of the drives gets narrower, so will the distribution of biases, with the distribution of the *s*_*i*_ preventing the distribution of biases from becoming arbitrarily narrow. For exponential𝒫(*s*), after many invasions the distribution of biases will approach a fixed width of order Σ and *L* will saturate at *L*_∞_ ~ 1/Σ^2^. But the mean *s* of the community, 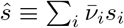, will steadily increase.

Successful invaders are unlikely to have *s*_*A*_ much smaller than *ŝ* (Section 4.5). Some will nevertheless have *s*_*A*_ < *ŝ*, but these can easily be driven extinct by later invasions. In contrast, invaders with *s*_*A*_ > *ŝ* will tend to survive for anomalously long (under successive invasions their bias is much less likely to decrease to *ξ*_*c*_). Thus on average invaders will increase *ŝ*. For the exponential distribution, once *ŝ* ≫ Σ, the lower boundary of the distribution no longer affects the distribution of *s*_*A*_ of successful invaders. Thus the distribution of *s*_*A*_ − *ŝ* will be independent of *ŝ*, due to the shift-invariance property of the exponential distribution. This distribution can be computed trivially. As the distribution of the drive, 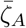, of potential invaders is gaussian and independent of *s*_*A*_, and their bias is 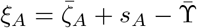, with the distribution of 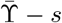 becoming independent of *ŝ*, the condition *ξ*_*A*_ > *ξ*_*c*_ for successful invaders enables integrations over the distribution.

The distribution of *s*_*A*_ − *ŝ* of successful invaders is proportional to 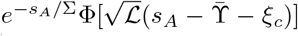 while the distribution of successful 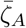 is proportional to 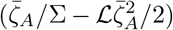 (see Section 6.10). Neither of these distributions has a lower limit but they decay as a gaussian for negative values and show the tilt to higher values due to the conditioning on successful invasion. In contrast, the distribution of biases, *ξ*_*A*_, of successful invaders is close to exponential, with scale Σ and lower cutoff at *ξ*_*c*_. Once *ŝ* has pushed into the tail of 𝒫(*s*), the convolution of the drive and *s* distributions will be dominated by the exponential tail of (*s*), and assumes a simple form as the lower cutoff of 𝒫(*s*) at 0 can then be neglected.

In the long-time steady-state for the exponential distribution, we have *L* ~ 1/Σ^2^ and the invaders have, on average, *s*_*A*_ − *ŝ* ~ Σ, thus the average increase in *ŝ* per successful invasion will be approximately constant: *dŝ*/*dZ* ≈ Σ/*L* ~ Σ^3^ (Figure 15). For a more detailed analysis, one could extend our Fokker Planck approximation in Section 6.8 to find the evolution of the joint distribution of 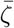 and *s*, and find a steady state solution which advances in 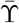 and *ŝ* at a constant rate. Similarly to the case without general fitnesses, *L* is fixed by the integral of the distribution and, by scaling, is proportional to 1/Σ^2^ (Section 6.9).

### 4.5 Diminishing-returns distributions of general fitness: *ψ* > 1

**Figure 9:**
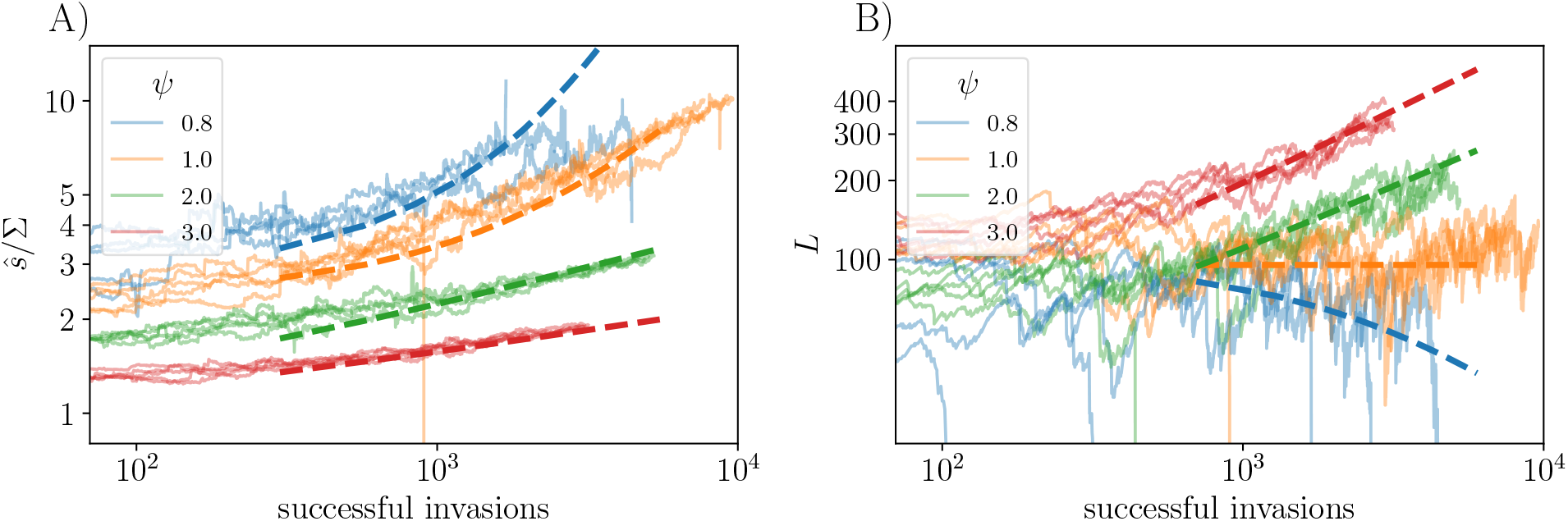
Comparison between predictions and scaling theory for unrelated invaders (*ρ* = 0), with general fitnesses, *s*_*i*_, drawn from distributions with tails parametrized by various values of *ψ*. (A) Increase of the community-average *ŝ* with successful invasions; 5 replicates of each simulation (solid lines) are shown along with theory curves (dotted lines). (B) Number of strains as a function of *Z* compared with theoretical predictions. In order to push up into the tails of the *s* distributions, the parameters of 𝒫(*s*) are different for the different values of *ψ*. The scales of the *s* distribution are here Σ ≃ 0.066, 0.08, .0177, .173 for *ψ* = 0.8, 1, 2, 3 respectively, and *K* = 300. The theory predictions are fit by eye, with forms 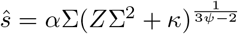, and 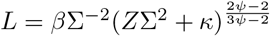. Values of *α, β* and *κ* for each *ψ* are reported in Table 6.11.

We now analyze the evolutionary dynamics for 𝒫(*s*) distributions that decay faster than exponentially: *ψ* > 1. As discussed earlier, if the *s* distribution of extant strains has pushed far enough out into the tail of 𝒫(*s*), then 𝒫 can be well approximated by an exponential distribution with scale 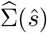 equal to the inverse log-slope of 𝒫(*s*) at *ŝ* as introduced in Equation 4. The above analysis of the exponential 𝒫(*s*) case is then valid until *ŝ* changes by enough that 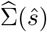 changes substantially. With 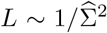, as expected, successful invaders will have 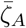 of order 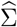 and 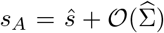 which, with *ψ* > 1, satisfies the criterion for being in the tail with 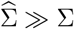. We thus expect that, after an initial evolutionary transient, the distributions of all quantities of the extant population and the successful invaders will be same as in the exponential case up to small corrections, with effective 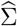 instead of Σ.

As evolution proceeds, *ŝ* slowly increases so that 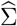 decreases and concomitantly *L* increases. From the above analysis, we expect 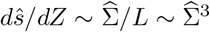. Integrating this equation and using 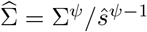 and 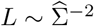, we can obtain scaling laws 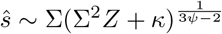 and 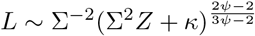 valid for *Z* > 1/Σ^2^. The effects of the order-unity coefficient *κ* roughly capture the transient regime with *L* ≪ Σ^−2^. For *ψ* > 1, we see that *dL*/*dZ* decreases steadily with *Z* since the behavior gets closer to an average of one extinction per successful invasion which obtains exactly in the steady state for the exponential distribution. In the initial transient, starting from *K* < 1/Σ^2^, the assembled community cannot yet feel the effects of the *s* distribution, so *L* will increase linearly in *Z*. In this regime we would expect roughly 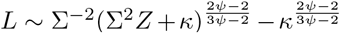. The initial rate of increase of *ŝ* will be slow since the effects of the *s*’s are small. The effect of *s*_*A*_ on the invasion probability of a strain is approximately linear in 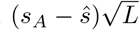 (normalizing by the width of the bias distribution). Since the *s*_*A*_’s are of order Σ, on average this results in an initial average increase in *ŝ* per invasion of 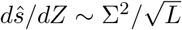.

The rate of successful invasions, *Z*, as a function of attempted invasions *Y* decreases as *ŝ* increases. In the independent invasion limit, it is given by the probability of drawing a general advantage greater than *ŝ* from the 𝒫(*s*) distribution:

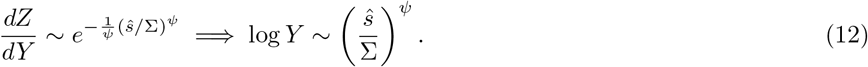

This scaling becomes valid once the distribution of *s* is pushed into the tail of 𝒫(*s*). In this regime, the probability of successful invasion decreases very fast with *ŝ*. Indeed, given the relationship between *ŝ* and *Z*, we have log *Y* ~ (Σ^2^*Z* + *κ*)^*θ*^ with 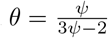 (see Figure 18). As the ecosystem evolves, it becomes harder to invade, with the total number of successful invasions,*Z*, only growing very slowly with the number of attempted invasions, *Y* (Figure 4). However, the number of strains versus *successful invasions* grows steadily for *ψ* > 1 in a way that is semi-quantitatively consistent with the predictions.

### 4.6 Correlated mutants without general fitness differences

We now give some preliminary and very partial analysis of the evolutionary behavior when the invaders are correlated with extant strains: specifically mutant-parent correlations of the interactions of magnitude parametrized by *ρ* < 1: E[*V*_*Mj*_*V*_*P* *j*_] = E[*V*_*jM*_*V*_*jP*_] = *ρ* with the newly-random parts of *V*_*Mj*_ and *V*_*jM*_ having variance 1 − *ρ*^2^ and correlation *γ*. To understand even a single mutant by dynamical mean field theory, one needs coupled equations to capture the interaction between parent, *P*, and mutant, *M*. Their direct interaction is small, ~ 1/*L*, as each has low abundance — but the effective interactions between them, mediated by interactions with the all the other strains, is comparable to their effective interactions with themselves 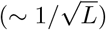. We have

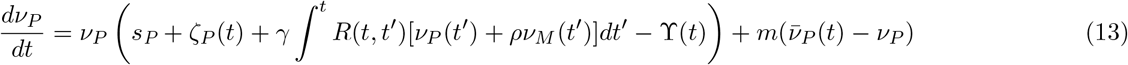

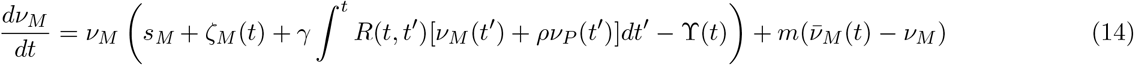

with the self-consistency condition on the correlations between the dynamic drives ⟨*ζ*_*M*_ (*t*)*ζ*_*P*_ (*t*′)⟩ = *ρ* ⟨*ζ*_*P*_ (*t*)*ζ*_*P*_ (*t*′)⟩ = *ρ* Σ_*j*_ ⟨*ν*_*j*_ (*t*)*ν*_*j*_ (*t*′)⟩. The feedback, *R*(*t, t*′), is the same sum of the responses over the other strains as in the uncorrelated case, but the feedback on the parent from the mutant history (and vice versa) is reduced by a factor of *ρ*.

The total time-averaged force from all the strains (neglecting migration) can, in the DMFT, be broken into several parts:

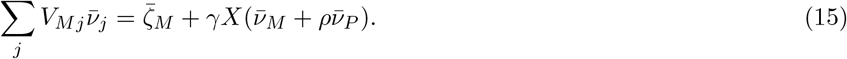

If both the parent and mutant have strongly positive biases in the absence of the other, then the effects of migration can — at least naively — be neglected and the DMFT equations for *d* log *ν*_*M,P*_ /*dt* can be averaged over time to yield

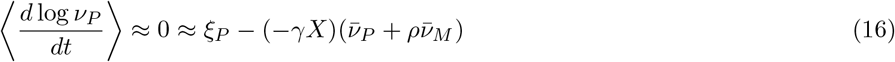

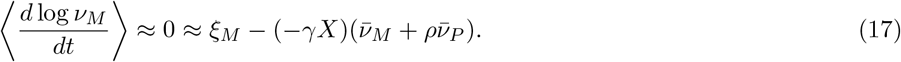

which is an effective two-strain competitive Lotka Volterra model from which the probability of coexistence between a parent and mutant can readily be found as done in Section 6.13. (Note that *γX* drops out of the expressions for coexistence and replacement probability, as long as it is negative as needed for a fixed point with positive abundances, and for the existence of the STC phase.)

**Figure 10:**
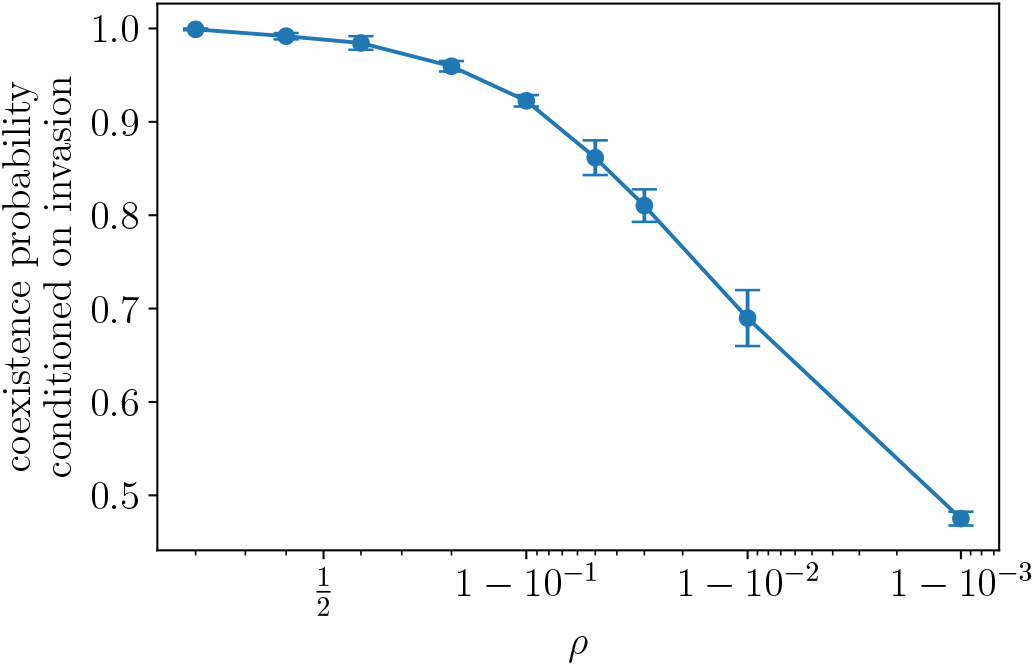
The coexistence probability between parent and mutant conditioned on successful invasion of the mutant (with all *s*_*i*_ = 0). The probability is averaged over 4 simulations of a diversifying community for each value of *ρ*, with error bars showing the standard error. Here *ρ* is shown on a logit scale to emphasize the difference between the points as *ρ* → 1. Even for *ρ* = 0.999, the coexistence probability is order 1/2 over the epoch of duration 30*ML*, long enough to enable small differences between closely related strains to have an effect.

Although the two-strain model would seem to be good for large positive bias — which parents are preferentially selected for as probability of a strain being chosen to mutate is 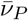 — it is certainly not good for smaller, or in particular negative, biases. Indeed, we find that the probability of coexistence between parent and mutant, conditioned on mutant invasion, is substantially larger than the simple two-strain Lotka Volterra equation would predict (Section 6.13). In the STC, a majority of persistent strains have negative bias and even for those with positive bias, the effect of migration (neglected in Equations 16 and 17) is not small. The bloom-bust dynamics with migration makes coexistence much easier. But understanding it even semi-quantitatively is challenging because of the correlations in the dynamical drive which makes parental blooms suppress the mutants and visa versa. Nevertheless, we expect that if *ρ* is extremely close to unity, the coexistence probability of parent and mutant will go to zero as 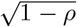 but with a small coefficient that might be of order some inverse power of *M* = log(1/*m*). In addition one should note that the natural timescale over which differences between the parent and mutant will accumulate is of order 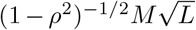 with the coefficient only ≅ 20 for *ρ* = 0.999. Therefore even for *ρ* this close to 1, the typical differences between parent and mutant will still be felt on timescales of order *ML* over which the close-to-marginal strains rise or die out.

The DMFT analysis of one parent and mutant, outlined above, becomes far more complicated once the community has evolved long enough that a substantial fraction of the strains have turned over (or coexist with relatives). The underlying simplification that makes DMFT valid for assembled communities, the approximate independence of the biases, will break down because of correlations among many of the strains due to their common ancestry. While some of the heuristic behavior of such communities and how they evolve may not change much even for *ρ* quite close to 1, there will certainly be quantitative changes.

The approximate Markovian analysis of the evolution of the bias distribution (Section 4.2), can be modified to roughly account for correlations between mutants and parents, *ρ* > 0. Such correlations will modify the effects of invading strains on the bias of the extant strains. This is most readily understood in the limit that *ρ* is very close to unity. In this case, one expects that the total abundance of the parent and mutant after the invasion will be similar to that of the parent before the invasion and, because of the high correlations, their effects on the other strains will be very similar. This suggests that both the stochastic and systematic changes of the biases should be multiplied by a factor of order 1 − *ρ*^2^ on the right hand side of Equation 10. For the first few mutants, the analysis could be carried through similarly. However once the number of successfully invaded mutants is comparable to the original *L*_0_, the correlations between the biases of the extant strains can not be ignored and the approximation of independent 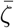’s breaks down. We leave analysis of this behavior for future work.

### 4.7 Correlated mutants with general fitness differences

**Figure 11:**
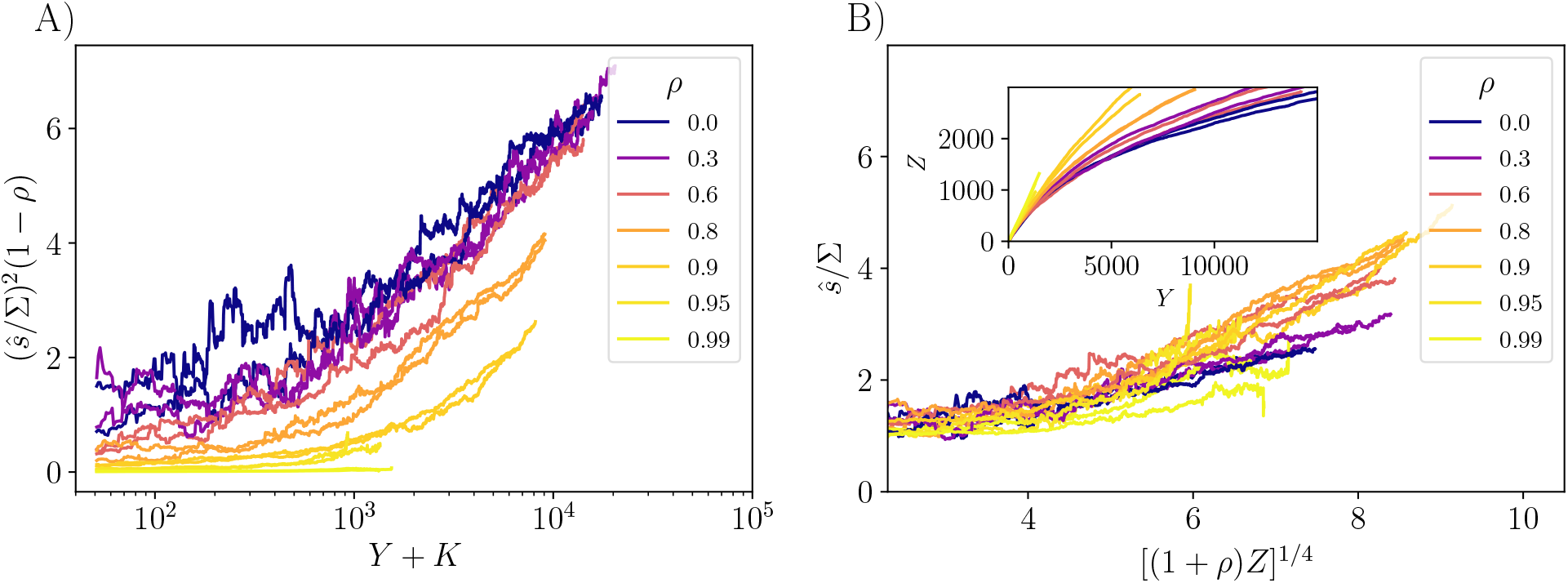
Evolution with correlated mutants and *s* distributed as a half-gaussian with standard deviation Σ ≅ 0.14. (A) The mean general fitness, *ŝ*, increases with attempted invasions for several values of mutant-parent correlation, *ρ*. Equation 20 predicts that 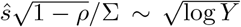, so on these axes, curves should approach straight lines with similar slopes. The offset of *Y* by *K* roughly accounts for the number of initial strains. (B) The same data as a function of the 1/4 power of the number of successful invasions, *Z*, rescaled by a factor of 1 + *ρ*, which although numerically small, is expected from Equation 19 and improves the comparisons. The theory predicts straight lines with similar slope across a range of *ρ*. Inset shows successful versus attempted invasions: Once evolution has pushed the *s*’s into the tail of 𝒫(*s*), the fraction of invasions that is successful is much higher for more correlated mutants since for small *ρ, s*_*M*_ is typically much less than *s*_*P*_ while for *ρ* → 1, *s*_*M*_ becomes close to the parental *s*_*P*_ and thus more likely successful.

The effects of correlated mutants with general fitness differences is complex. We see that correlations tend to speed up the rate of evolution because they draw *ss*’s of new strains from a distribution with higher mean than in the uncorrelated case. This increases both the fraction of successful invasions, and the increases in *ŝ* associated with successful invasions.

For general *ψ*, we draw the mutant *s*_*M*_ from a distribution conditioned on *s*_*P*_ that preserves the tail of 𝒫(*s*):

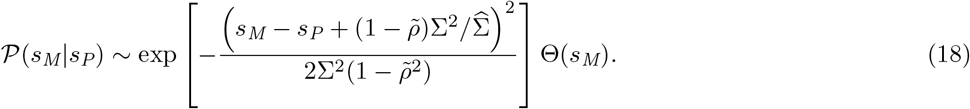

Note that this expression reduces to Equation 5 when we set = 2, which results in 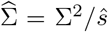. In that gaussian case, we also preserve the conditional mean E[*s*_*M*_|*s*_*P*_] = *ρs*_*P*_ (plus corrections due to the truncation of the distribution). For convenience, we set the correlation parameter for the *s*’s to be the same as for the interactions: 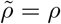 (as in simulations), but having these parameters unequal is a relevant extension that we have not studied.

Since we preferentially choose successful parents (with probability proportional to their average abundance), we are likely to have *s*_*P*_ ≅ *ŝ*. In the *ψ* = 2 model, the local slope of the log-distribution of the mutant *s* evaluated at distance (1 − *ρ*)*ŝ* from the the mean at *ρŝ* is 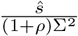, and the number of strains scales as 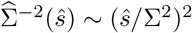. The rate of increase of the community-average *ŝ* with successful invasions is

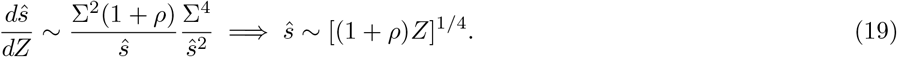

Therefore we expect a weak positive dependence of *dŝ*/*dZ* on *ρ*, consistent with Figure 6. From the shape of the conditional distribution of the mutant’s general advantage, we expect that

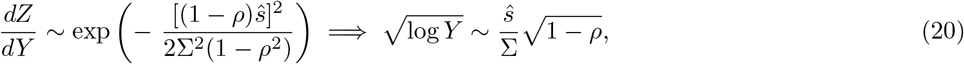

where we omit factors of 1 + *ρ* which do not affect the scaling as *ρ* → 1.

When *ρ* → 1, there are various subtleties discussed above and below which make comparing simulations and theory difficult. The timescale for highly correlated parents and mutants to drive each other extinct becomes very long, and potentially comparable to the timescale from stochastic finite-island-number extinctions. Furthermore, the nature of the chaotic phase seems to stabilize coexistence between very closely related strains to an unexpected degree, as discussed above. We leave further exploration of the interplay between closely related strains for future work.

With correlated general fitnesses, as *ŝ* increases there is a crossover scale of *L* beyond which it becomes much harder to push into the tail of 𝒫(*s*). Looking at 𝒫(*s*_*M*_ ≥ *s*_*P*_ |*s*_*P*_) as a rough estimate of the probability of invasion (which its dominated by *s*_*P*_ ≈ *s*_*M*_) suggests that if 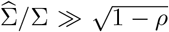 then it is relatively easy to invade — the mutant strain has a substantial chance of drawing *s*_*M*_ at least as big as *s*_*P*_. However if 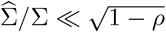 then it becomes very unlikely for *s*_*M*_ to be larger than *s*_*P*_. In the exponential case, 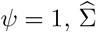 and *ρ* are constants so their values determine the regime of evolution — but for *ψ* > 1 there will be a change in behavior as *ŝ* increases, and after the crossover the probability of successful invasion becomes gaussian small in 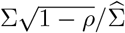. The value of *L* at this crossover is Σ^−2^(1 − *ρ*)^−1^, which diverges as *ρ* → 1. Therefore most of our simulations remain in the regime before the crossover.

### 4.8 Abundance distributions and interactions conditioned by evolution

A crucial question is how the properties of evolved communities differ from communities assembled from separately evolved strains. The simplest observable properties are distributions of abundances: either the roughly-equivalent time- or space-averaged abundances; or a *snapshot* of abundances at one location and one time. We can compare these distributions between two similar size communities: one that diversified from a small number of initial strains, and the other an unevolved assembled community persisting out of a larger number of initial strains.

**Figure 12:**
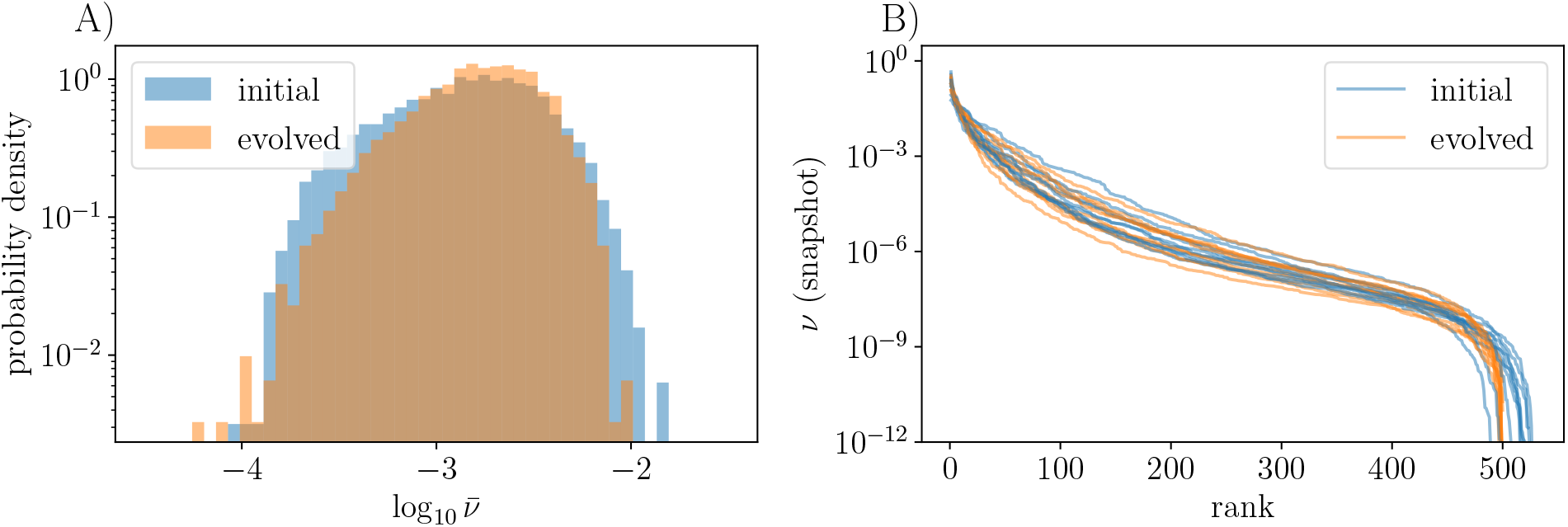
(A) Distribution of mean abundances for assembled and evolved communities (of unrelated strains with no general fitness differences). Communities have *L* ≈ 500, where evolved communities have diversified from *K* = 50 initial strains so that they lose memory of their initial assembly conditions, and assembled communities have *K* = 650 with *L* = *L*_0_ ≈ 500 surviving after the long epoch used. Though the bulk of the distributions looks similar, there is a marked depletion in the likelihood of very rare or very abundant strains in the evolved community. (B) Rank abundance distributions from snapshots on a single island (each curve is from a different simulation) look quite similar between assembled and evolved communities. The evolved community shows fewer strains at low abundance which is also evident in the average strain abundances from panel (A).

The mean abundances in an evolved community are somewhat more narrowly distributed than in an assembled community, with fewer highly abundant and fewer rare strains. This is consistent with our picture of the bias distribution being smoothed out at the low end due to invasion-triggered extinctions: the depletion of low bias strains results in fewer strains with low abundance. The depletion of abundant strains is likely due to the kill-the-winner dynamics which rewards invading strains that push the must abundant extant strains down. For the snapshot distributions, we plot the rank-abundance distribution on one island at an instant in time; this shows a similar modest depletion of low abundance strains in the evolved communities as compared to the assembled communities. [Note that most widely-used measures of diversity are not really informative for these kinds of distributions that are broad on a logarithmic scale. For example the Shannon entropy would weight mostly the highly abundant strains, while the “species richness” would be highly sensitive to the cutoff in observable abundance.]

Another aspect to compare, is the distributions of the interactions between extant strains. How do the statistics of the interaction matrix _*ij*_ change when conditioned on non-extinction in an assembled community? When conditioned on evolutionary history? Running a set of 10 replicates without general fitness differences over a period where the community grows from 50 initial strains to ≅ 500 strains for both *ρ* = 0 and *ρ* = 0.95, we obtain ensembles of interaction matrices conditioned on gradual evolution. We also examined the interaction matrices of communities that evolved at steady state in *L* — using an exponential distribution of *s* with Σ = 0.07, starting from *K* = 50 and reaching steady state with *L* ≅ 150 while evolving over the course of ≅ 4000 successful invasions. In this case one might be more likely to see correlations building up, since new strains might not as effectively dilute the correlations building up in *V*. For contrast, we study assembled, (but not evolved) communities starting with sufficiently large *K* that *L*_0_ ≅ 500. The statistics of the conditioned *V*_*ij*_ are compared with those of random sampling from the unconditioned distribution.

In Table 2 we define a number of statistics which one might expect to change over the course of the evolution. These include 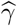, the empirical cross-diagonal correlation in the *V* matrix of extant strains; 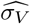, the empirical width of the interaction strength distribution after evolution; 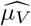, the average of the interactions; and 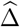, the standard deviation of the distribution (over the *i*) of Σ_*j*_ *V*_*ij*_, which is like an effective 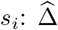 crudely approximates the width of the bias distribution. Aside from 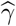, all the quantities are defined including the diagonal terms in *V*.

**Table 1:** Statistical properties of the interaction matrix for evolved and assembled communities: 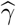 is the cross diagonal correlation in *V*, and 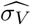 the width of the *V*_*ij*_ distribution; 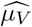 is the mean of the *V*_*ij*_ and 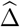 is the standard deviation (over *i*) of Σ_*j*_*V*_*ij*_, which is like an effective *s*_*i*_. Evolved communities grew to ≅ 500 strains from an initial *K* = 50 and assembled communities had *L*_0_ ≅ 500 after a single epoch of duration 12*ML*, both with all *s*_*i*_ = 0. The last line is from a simulation with an exponential distribution of *s* and Σ = 0.07, which results in about 150 strains at steady state. The mean of each quantity is calculated across 10 replicates. For comparison, the approximate statistics from the original ensemble (“pool”) with *K* = 500, *γ* = −0.8 are given (modulo slight corrections arising from the diagonal terms). In Table 5 we show the statistics of the conditional *V* normalized by the standard deviation of the matrix *V* from the pool, which allows one to determine which changes are statistically significant. Purple entries correspond to those more than 3 standard deviations away from the pool value.

| | $\widehat{\gamma}$ | $\widehat{\sigma}_V$ | $\widehat{\mu}_V$ | $\widehat{\Delta}$ |
| --- | --- | --- | --- | --- |
| pool ($K = 500, \gamma = -0.8$) | -0.8 | 1 | 0 | 0.0446 |
| assembled ($\Sigma = 0$) | -0.8014 | 1.0012 | 0.0026 | **0.0337** |
| evolved ($\Sigma = 0, \rho = 0$) | -0.8015 | 1.0028 | **0.0061** | **0.0289** |
| evolved ($\Sigma = 0, \rho = 0.95$) | **-0.8082** | **1.0427** | **0.1422** | **0.0328** |
| pool ($K = 150, \gamma = -0.8$) | -0.8 | 1 | 0 | 0.0816 |
| evolved ($\Sigma = 0.07, \rho = 0$) | -0.8074 | **1.0205** | **0.0165** | **0.0977** |

**Table 2:** Definition of some statistics of the interaction matrix of a community.

| $\hat{\gamma}$ | $\hat{\sigma}_V$ | $\hat{\mu}_V$ | $\hat{\Delta}$ |
| --- | --- | --- | --- |
| $\text{Corr}[V_{ij}, V_{ji}], i \neq j$ | $\text{Stdev}_{ij}[V_{ij}]$ | $\frac{1}{L^2} \sum_{i,j} V_{ij}$ | $\text{Stdev}_j [\frac{1}{L} \sum_i V_{ij}]$ |

**Table 3:**
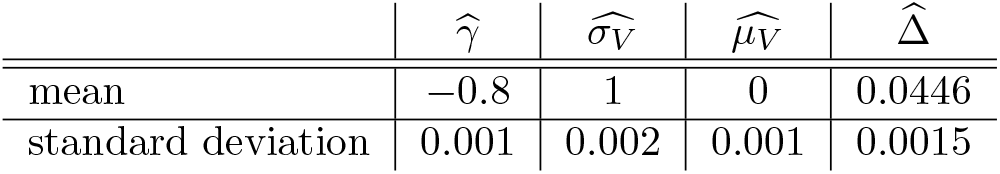
We can estimate the standard deviation of these quantities from an ensemble of 500 realizations of the matrix *V* with *γ* = −0.8, *L* = 500. For 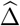 we also estimate the mean from the numerics.

| | $\hat{\gamma}$ | $\hat{\sigma}_V$ | $\hat{\mu}_V$ | $\hat{\Delta}$ |
| --- | --- | --- | --- | --- |
| mean | -0.8 | 1 | 0 | 0.0446 |
| standard deviation | 0.001 | 0.002 | 0.001 | 0.0015 |

**Table 4:**
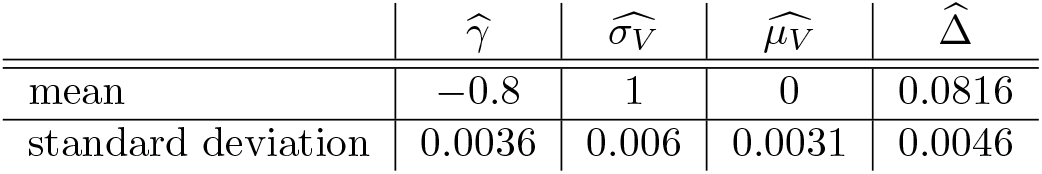
We also calculate the mean and standard deviation of these quantities over 500 matrices for *L* = 150 and *γ* = 0.8, which is pertinent for analyzing results from evolution at constant *L* ≅150 set by an exponential 𝒫 (*s*) with Σ = 0.07.

| | $\hat{\gamma}$ | $\hat{\sigma}_V$ | $\hat{\mu}_V$ | $\hat{\Delta}$ |
| --- | --- | --- | --- | --- |
| mean | -0.8 | 1 | 0 | 0.0816 |
| standard deviation | 0.0036 | 0.006 | 0.0031 | 0.0046 |

**Table 5:** Properties of the interaction matrix for evolved and assembled communities, displayed in terms of number of standard deviations from the mean in the original *V* ensemble. These are the same data as in Table 1, but normalized according to appropriate scale of deviations calculated from original *V* ensemble without conditioning. Numbers in purple have magnitude greater than 3, and are thus clearly statistically significant.

| | $\tilde{\gamma}$ | $\tilde{\sigma}_V$ | $\tilde{\mu}_V$ | $\tilde{\Delta}$ |
| --- | --- | --- | --- | --- |
| assembled | -1.4 | 0.6 | 2.6 | -7.2 |
| evolved (diversifying), $\rho = 0$ | -1.5 | 1.4 | 6.1 | -10.4 |
| evolved, $\rho = 0.95$ | -8.2 | 21.4 | 142.2 | -7.9 |
| evolved (steady state) | -2.1 | 3.4 | 5.3 | 3.5 |

Our results show that both evolution and assembly tend to make 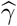 slightly more negative than the *γ* of the strain pool, and that these processes also favor an increasing mean interaction strength, 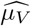. Of particular interest, given the previous discussion, is 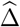. For *L* ≅ 500, the statistics of the matrices from the pool have 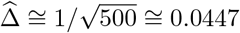 However once we condition on assembly or evolution, the value of 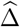 can be expected to decrease, because we eliminate those strains with negative bias, meaning that strains *i* with an anomalously small value of Σ_*j*_ *V*_*ij*_ will go extinct, making the range of Σ_*j*_ *V*_*ij*_ narrower. This is indeed observed in Table 1, at least for simulations without general fitness differences. Interestingly we see that 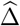 decreases with evolution by more for *ρ* = 0 than for *ρ* = 0.95, which could be because, in the latter case, the elements in a single row of *V* are correlated with those of a high-abundance parent, which could increase the width of the Σ_*j*_*V*_*ij*_ distribution. Nonetheless we still expect that 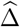 scales as 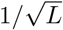 for evolved communities.

For the evolution with *L* roughly constant from the pure exponential distribution of *s* (*ψ* = 1) with Σ = 0.07, we see that 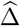 after evolution is actually larger than in an unconditioned matrix (see the last two rows of Table 1). This could be due to the buildup of correlations in the interactions between extant strains.

We can compare these results to previous results of Bunin [28] in which he calculated the statistics of the interaction matrix conditioned on assembly in the stable fixed-point phase where niche interactions rather than spatial structure stabilize the diversity. Bunin observed that, conditional on assembly, correlations between the effect of strains *k* and *l* both on strain *i* become negative, i.e. Corr[*V*_*ik*_*, V*_*il*_] < 0 with small magnitude of order 1/*K*. This is consistent with our observation that the distribution of the sum of these interactions for each strain, Σ_*j*_ *V*_*ij*_, will have a smaller width than in the pool. In addition, Bunin finds an 𝒪(1/*K*) shift in the mean of the interaction matrix toward less competition, which is consistent with 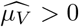. We also see slight but systematic changes in 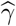 and 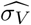, particularly in the case of evolution of correlated mutants with *ρ* = 0.95.

Although many of the changes of the interaction statistics of assembled or evolved communities are statistically significant (Section 6.12), they are mostly very small, with the possible exception of 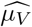 for *ρ* = 0.95. We conjecture that for *ρ* = 0 all the effects are small by some power of *L*. But for *ρ* near one, this is less clear: whether for *L* large compared to some inverse power of 1 − *ρ*, the correlations induced by evolution will still be small, or whether they will persist for arbitrarily large *L*, we leave as an open question.

### 4.9 Nucleation from low diversity

**Figure 13:**
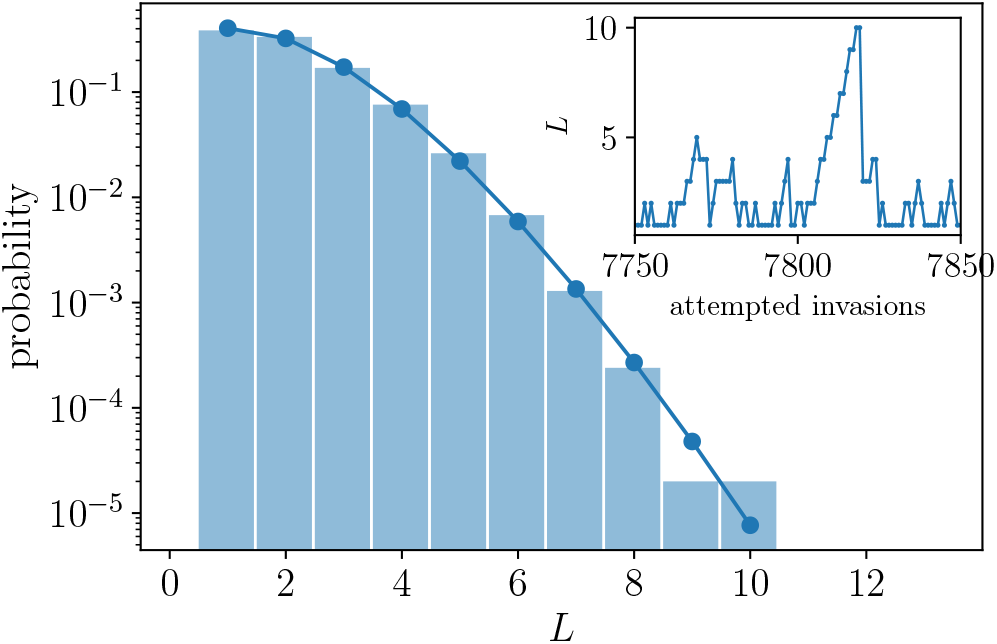
Distribution of community size *L* starting from a single strain, over 10^5^ attempted invasions. The solid line shows a Poisson fit with mean 1.6, excluding *L* = 0. This fit is remarkably good. The inset shows the trajectory of *L* as it reached its maximum value which occurred only once, followed by a crash back down.

We conclude this analysis section by further investigation of the evolutionary dynamics when the diversity is low. We are particularly interested in the nucleation process by which an ecosystem could transition from the *L* ≪m *L*^*^ regime to the steadily diversifying regime. For the parameters we have investigated, this turns out to be extremely rare. Even in runs of 10^5^ attempted invasions starting from a single strain, the community always remains small. We observe that the number of extinctions per successful invasion is broadly distributed (as when starting with a larger community *L* ~ *L*^*^ in Figure 3) and the evolutionary dynamics of *L* typically proceeds by incremental increases punctuated by large decreases. In this regime, the steady state distribution of *L* is observed to be very close to Poisson (Figure 13) — a result that we do not have a theory for. From this Poisson behavior, one can attempt to extrapolate the probability of reaching *L*^*^ and transitioning to steady diversification by a lucky fluctuation in *L*. Taking *L*^*^ ≈ *L*_0_(*K*^*^ ≈ 70) ≈ 55 based on Figures 16 and 3, this estimate suggests ~ 10^60^ attempted invasions would be needed. However the probability distribution of *L* will surely deviate from Poisson long before *L*^*^, so our prediction for the chance of nucleation is likely too small by many orders of magnitude, depending on the value of *L* for which the probability distribution starts to be significantly higher than the Poisson extrapolation. At this point we do not have even conjectures for the shape of this distribution, nor for how the value of *L*^*^ changes with *γ, ρ* or *m*. At the marginal point *γ* = − 1, we have the special ASM, and diversification becomes impossible, so near there we expect that *L*^*^, along with the fragility, diverges.

A more in-depth analysis of the nucleation process should consider the mechanism of synchronization [53], or more relevantly, *desynchronization* which is necessary to enter the steadily diversifying regime. In fact we observed that decreasing the migration rate *m* increases the mean of the Poisson-like distribution of *L* in the low-diversity regime, and also reduces *L*^*^ (Figure 16), thereby perhaps substantially increasing the probability of nucleation — although still too low to observe.

## 5 Discussion and Questions

In this paper we have answered an important issue of principle: Without any assumption of niche-like differences between strains, can the diversity continually grow under slow evolution? We have found that this can indeed occur if the community forms a spatiotemporally chaotic phase that we have studied previously in PAF. As new strains are introduced — either separately evolved invaders or mutants of extant strains — some successfully invade, and some of these drive extinctions of one or a few of the extant strains. In a range of parameters, the size of the persistent community continually grows on average, while for other parameters, the diversity shrinks and eventually crashes. How fast the diversification proceeds depends on the statistical properties of the strains. If each strain has different general, nonspecific fitness, *s*, then as evolution proceeds the average *s* of the population gradually increases and pushes into the tail of the *s* distribution. If the high-*s* tail falls off faster than exponentially, the community continues to diversify but more and more slowly, since fewer new strains will have sufficiently large *s* to invade. For broader-than-exponential distributions, the diversity eventually crashes as the general fitness differences dominate over the the effects of interactions with other strains.

Building on analytic and scaling understanding of the STC phase for an assembled community, we have developed a substantial understanding of the dynamics of the diversification or de-diversification. However even for the simple models on which we have focused, there are aspects that we do not understand.

### 5.1 Unresolved issues with the simple island models

#### Development of correlations

Even with invaders uncorrelated with the extant strains, subtle correlations build up in the interaction matrix and — although they appear rather weak (Section 4.8) — the memory of earlier evolution will affect the way strain abundances change under further evolution, mandating a better treatment of the evolution than the Markovian approximation we have used in Section 4.2. With several complicating features — mutants, correlated nonspecific differences, and substantial-sized initial communities — included, there are a number of crossovers that we have not attempted to analyze (see Section 4.7). These, and which aspects promote, slow down, or prevent, continual diversification, are likely to be quantitative and strongly model-dependent.

#### Nucleation of diversifying “phase”

An observation from the simulations (Section 3.1) gives rise to a broader question: Why is it so hard to nucleate the diverse STC phase? And, concomitantly, why do initially diverse communities so often crash unless the diversity is rather large? It is likely that the limited number of strains that dominate on each island over any short time interval — of order *L*/ log(1/*m*) — plays a role, but unclear how. Whether the difficulty of nucleating a diverse STC community is special to the structure of the models and spatial dynamics assumed, or is true more generally, certainly needs further investigation.

#### Spectrum of mutants and coexistence of parents and mutants

When invaders are mutants of extant strains that differ from their parent only very slightly, (with correlation coefficient *ρ* very close to unity), we have found that the parent and mutant coexist surprisingly frequently. Understanding this, even for the first mutant, requires analyzing the dynamics of strains with strongly correlated noise which we have not carried out, although we suspect that the very large local abundance variations that occur with low-migration rate give rise to a small decorrelation scale needed for coexistence. In each simulation, we have considered only mutants with a fixed level of correlation with their parents, leaving a number of natural questions: What are the effects of a distribution of magnitudes of mutational differences? How do these affect the invasion, coexistence, and subsequent properties of the evolving communities?

#### Invasion dynamics

Because of the local chaos and low migration, the invasion of a potentially-successful new strain is complex. To avoid this complication, we have introduced new strains at substantial abundance and on all islands simultaneously. In actuality, most initial invasion attempts on an island will fail: only if the strain arrives when the conditions are ripe for it to bloom, can it avoid quick extinction and send out enough offspring to other islands, which — if also sufficiently good timing — allow it to spread. How this process depends on the relatedness of mutant and parent complicates matters greatly because of the boom-bust dynamics. Strains are most likely to beget mutant offspring when their abundances are high, but at that stage of a bloom, a crash in the local population will soon follow. Though many mutants may arise when a parent strain is doing well, the correlation between their dynamics and those of their parent means that they are likely to quickly go extinct when their parent population crashes down from high abundance. In contrast, mutants that emerge right before a parent blooms from low abundance up to high abundance, can ride the bloom and establish more readily, but they would have to arise in a small parental population. Understanding the balance between these effects and their consequence for invasion probabilities is a challenge for future work — especially with real spatial structure and dynamics, discussed below.

### 5.2 Phylogenies and effects of correlations between strains

Once multiple mutants have established in a community, given rise to further descendants, and so on, the potentially-hierarchical correlations between the dynamics of different strains will make the correlated dynamics complex and any analysis a major challenge: reliance on simulations for observations and to frame conjectures is needed. For this, focusing initially on models with the least complicating features may be advisable.

The most readily observable features of the relatedness in an asexual community are the phylogenies. Are the statistical properties of the phylogenies — here driven entirely by “selection” in the broad sense, but with the ecological interactions creating a balance between the many extant strains — similar to a known class of coalescent trees? Simulating many large enough communities may be hard. But if neutral mutations could be efficiently included, in addition to the phenotype-changing mutations studied thus far, more data could be taken. Would the distinctions between the neutral and non-neutral mutations be apparent?

An important quantitative characteristic associated with the hierarchical relatedness in an evolved community is the distribution of the biases. The reason that a distribution of general fitness differences limits diversification is that the variations in the biases across the community decreases as 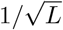 and thus is easily dominated by nonspecific differences. Can hierarchical correlations make the bias distribution broader? In principle, if the differences between the most divergent strains remained large enough as the community continued to diversify, the variations in *s* might not dominate and “entropy” might favor the mean *s* saturating rather than creeping further into the tail of the *s* distribution. If so, perhaps the diversification could continue much more rapidly. In our preliminary investigations we have not seen evidence of this occurring — even with *ρ* = 0.95 the simple correlations that evolve are modest — but we have not ruled it out: the important correlations might be much more subtle.

### 5.3 Spatial structure and dynamics

While for some microbial populations — for example common human gut commensals — a collection of connected “islands” without much spatial structure may be a rough caricature, in most there is spatial structure that makes dispersal from one location to another dependent on the distance in one, two or three dimensions. Thus instead of having all pairs of islands connected by migration, one could model a *d*-dimensional array of islands with nearest-neighbor migration; a spatial continuum with diffusive dispersal; or a mixture of long and short distance dispersal events as driven by wind, ocean currents, or hitchhiking on migrant animals [54]. With real spatial structure, local sub-populations are much more prone to extinction and cannot be as readily rescued by migration from another location where the strain is blooming. Thus in contrast to the regime we have worked in for this paper, recovery from local extinctions must play a crucial role. The dynamics of invasions, extinctions, and repopulation is very different than in the spatial mean field model: if the underlying dynamics is diffusive, invasion and repopulation will occur by propagating Fisher-Kolmogorov-Petrovsky-Piscunov (FKPP) fronts [46]. The properties of FKPP waves are known to be highly sensitive to dynamics at the wavefront, and the effects of demographic fluctuations have been investigated [47]. But the approximately multiplicative “noise” from the ecological interactions will surely change this, and even for a single wave understanding the impact of these larger fluctuations is still an open question [55, 56].

With long-range dispersal over a multitude of length scales, the dynamics of invasion, extinction and repopulation will be very different, as already occurs for a single successful invader without ecological variations [54]. Generally, understanding of the STC phase will have to build on better understanding of repopulation dynamics in the presence of large ecological fluctuations, and then understanding the evolution of communities on top of that. We leave investigations of this for future work. But we conjecture that a continually diversifying STC phase can still occur with real spatial dynamics.

### 5.4 Bacteria-phage interactions and coevolution

An obvious weakness of the Lotka-Volterra models studied here is that the strains do not carry their own phenotypes, but are characterized by their interaction with all possible other strains. Furthermore, the antisymmetric correlations in the interaction matrix (especially without substantial nonspecific differences) are rather unnatural for multiple strains of a single species. Thus the most interesting extension of this work is to much more natural models: multiple strains of a phage species that prey on multiple strains of a bacterial species, with varying effectiveness that is a function of phenotypic properties of the particular phage and bacterial strains — in particular of the interaction between a phage tail and bacterial receptor. We showed previously (in PAF) that the block-antisymmetrically-correlated structure of the interaction matrix with the bacteria having no niche-structure (differing only in the way they interact with the phages) can give rise to an STC phase that is very similar to that of the antisymmetrically correlated Lotka-Volterra model studied here. Such a bacteria-phage model naturally includes the possibility of mutants having general fitness advantages, obviating the need to introduce the nonspecific differences on a separate footing. In ongoing work, we show that much of the basic phenomenology we have found here also occurs in evolving bacteria-phage phenotype models — at this stage only roughly and qualitatively. In this context, the phylogenies and relatedness questions are particularly natural. Whether more specialist phages tend to evolve, making the interaction matrix sparser and perhaps more hierarchical — and if so under what circumstances — is also a natural question.

### 5.5 Concluding questions

We have studied evolution of communities of many closely related strains in the limit that the evolutionary dynamics is slow compared to ecological and spatial dynamics. For a class of models, and in a particular ecological “phase,” evolution drives continual diversification, provided there is sufficient diversity initially. However mutations that change general fitness tend to strongly slow down or even reverse the diversification. How ubiquitous is such diversification in the absence of any niche-like structure? Are there models in which a diversifying phase is easier to nucleate? Will the diversification always tend to be limited or strongly-slowed by nonspecific mutational effects? Or might “entropic”-like effects of difficulty finding such mutations — e.g. from discrete genomes rather than continuous phenotypic parameters, or from soft tradeoffs — counter this, or perhaps produce evolutionary dynamics that lead to sparse interaction matrices and broader distributions of biases? Conversely, if strains are initially separated in “niche space” but then start to overlap and interact as the number of strains increases, how does the behavior differ? Is continual diversification easier to nucleate?

What happens when, as in large microbial populations, evolutionary processes are not slow? Faster evolution is likely to make diversification easier, but understanding this even in simple models will require much better understanding of the invasion probabilities of mutants. Other than our scenario in which spatiotemporal chaos is the key to stabilizing coexisting diversity, what other robust scenarios are there? And of course, most crucially, what observable features of the strain, sub-strain, and sub-sub-strain level diversity in a microbial population (or interacting populations) could provide hints to the underlying causes of extensive diversity?

## Acknowledgements

This work was supported in part by the National Science Foundation via PHY-1607606, the National Institutes of Health via R01AI13699201, and the Stanford University Sherlock Cluster for the computations.

## 6 Appendix

### 6.1 Numerical implementation

Parameter values unless otherwise mentioned are *I* = 40, *m* = 10^−5^, *γ* = −0.8, *N* = 10^20^.

For numerical integration we use an adaptive forward Euler step, with the time step chosen so that the maximum fractional change in the abundance of any strain is no greater than 3/4. In other words, in a single time step an abundance can change to anywhere from 1/4 to 7/4 of its current value. Because of the wide range over which abundances vary, such large changes do not cause numerical problems.

To save computational time, we reject some incoming strains because they are very unlikely to successfully invade. This can be estimated by calculating their bias from the properties of the community before their introduction. By roughly estimating the critical bias from earlier strain invasion attempts, we can conservatively reject some incoming strains without having to run the dynamics — with nonspecific differences this can yield substantial speed-up by rejecting strains with *s* − *ŝ* sufficiently negative.

### 6.2 Epoch duration

Upon initializing our simulations, we allow an ensemble of *K* randomly-drawn strains to relax under the dynamics of Equation 1 for time 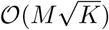. Between the introduction of new strains we run the ecological dynamics for epochs of length *c*_epoch_*ML* with *c*_epoch_ = 3. In addition, for most simulations we intersperse regular-length epochs with longer epochs to allow the ecosystem time to reach steady state. We inserted longer epochs (longer than the others by a factor of ~ 3) every 100 epochs to allow marginal strains to go extinct.

The duration of the initial epoch after a community of *K* strains was assembled, was usually only 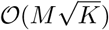 (which does not much matter as marginal strains can in any case go extinct in subsequent epochs). However, for the simulations where we compared assembled and evolved communities (e.g. Figure 12), the dynamics of the assembled communities were run for time 12*ML*.

**Figure 14:**
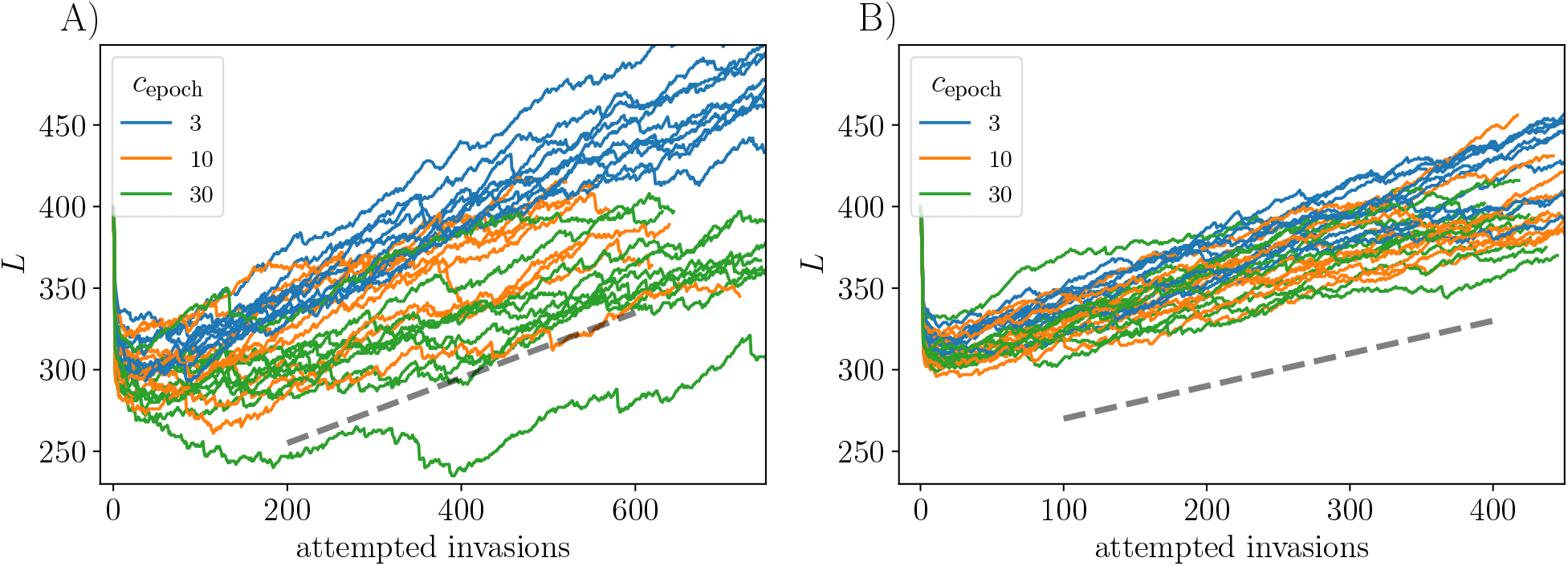
*K* = 400. (A) *ρ* = 0. (B) *ρ* = 0.95. Rates of diversification (measured per attempted invasion) are insensitive to a factor-of-10 change in the intervals between the introduction of the new types: the epoch durations are *c*_epoch_*ML*. Although it can take more invasions to get into the steadily diversifying regime for long epochs, the rates of diversity increase depend little on the epoch lengths. For this comparison, it also happens that the rates are similar for *ρ* = 0 and *ρ* = 0.95: dashed lines both have slope 0.2.

In order to check that the evolutionary dynamics is in the quasistatic limit, where the strains are introduced slowly enough that their introduction rate does not matter, we look at how the diversification rate depends on the length of the epochs between strain introductions. In Figure 14 we vary the *c*_epoch_ over one order of magnitude and see similar dynamics for all epoch lengths. In the case of *ρ* = 0, there is a longer transient for the long epochs, though once in the diversifying regime, *L* increases at a similar rate for all epoch lengths. Based on this, all data presented in the paper are generated with *c*_epoch_ = 3, which makes numerics faster, and should not change qualitative conclusions about the slowly diversifying regime. Though there is a slight slowdown in diversification for longer epochs, keeping the just-below-marginal strains which would go extinct in the longer epochs, does not affect our conclusions: in any case these marginal strains contribute little to the dynamics of the community. For *ρ* = 0.95 there appear to be few marginal strains, possibly because preferentially mutating the strains with large abundance is unlikely to produce descendants with close-to-marginal bias.

### 6.3 Correlations between mutant and parent interactions

The effects on a strain of a mutation is comprised of two parts: changes in its *interactions* with the other strains, and a change in its general fitness *s* yielding a systematic change in its growth rate. To generate the interaction part of a mutation, we append a new row and column to *V* which parameterizes the interaction between the new strain *M* and the parent strain *P*, each with another strain, *k*, according to E[*V*_*Mk*_*V*_*P* *k*_] = E[*V*_*kP*_*V*_*kM*_] = *ρ* for *M, P* ≠ *k*. The parts of the mutant’s interactions, *V*_*Mk*_ and *V*_*kM*_, that are not correlated with the parent have correlation *γ* between them, so we take 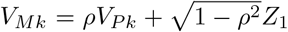 and 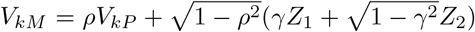 where the *Z*_*i*_ are i.i.d. standard normal random variables. This preserves the desired correlations, with E[*V*_*Mk*_*V*_*kM*_] = *γ* if the expectation is also over unevolved parental interactions.

However we have to treat the direct interactions between the parent and mutant, *k* = *M, P*, more carefully. Define *H* as the effect of the mutant on the parent, *J* as the effect of the parent on the mutant, and *D* and *K*, the respective self interaction terms, i.e.

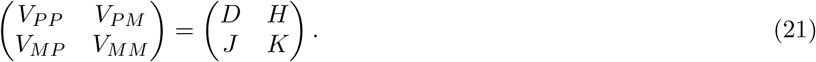

The symmetry conditions we want to enforce are

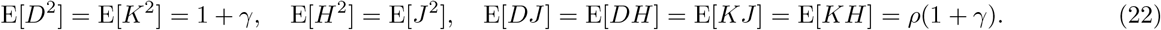

In addition we require that *D* = *H* = *J* = *K* in the limit *ρ* → 1, and E[*HJ*] = *γ* as *ρ* → 0. The parameterization that we choose which respects these conditions is

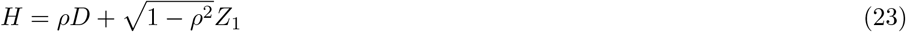

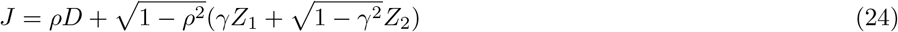

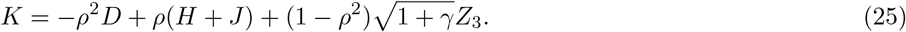

where the *Z*_*i*_ are i.i.d. standard normal random variables. Now we have E[*HJ*] = *ρ*^2^ + *γ*(1 − *ρ*^2^), which does not precisely preserve the desired correlation E[*HJ*] = *γ*, but it has the correct limit as *ρ* → 0. Our parameterization choice gives E[*DK*] = *ρ*^2^(1 + *γ*). There is some arbitrariness in this parameterization as it is possible to choose an alternative which has a different value of E[*DK*] but still respects the desired symmetries.

The dominant effective interaction between the parent and the mutant is mediated by the other strains in the ecosystem as in the analysis from Section 4.6. The direct influence of the parent on the growth rate of the mutant and vice versa is smaller than the contributions mediated by all the other strains by a factor of 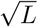, thus the choice of the direct parent-mutant interaction ought not to be of much importance — unless sufficiently strong correlations develop under continuing evolution, for which we do not see evidence.

### 6.4 *dŝ*/*dZ* in the simple exponential (*ψ* = 1) model

**Figure 15:**
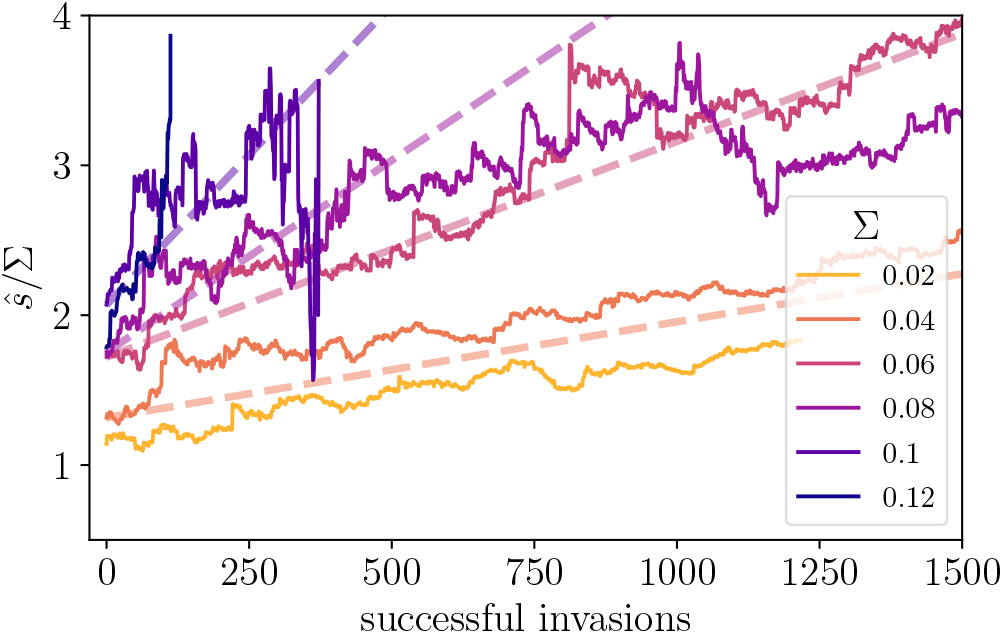
Community-mean general fitness, *ŝ*, as function of number of successful invasions, *Z*. From the scaling argument *dŝ*/*dZ* ~ Σ/*L*, we expect that *dŝ*/*dZ* ~ Σ^3^. For the runs in which Σ is high enough that the diversity crashes, we nevertheless observe the expected scaling momentarily before the crash. The rate of *ŝ* increase per successful invasion scales with Σ^3^, so normalizing by Σ we have 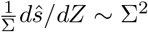. Here we plot *ŝ*/Σ versus successful invasions; the dashed lines have slope 0.4 × Σ^2^ for the intermediate range of Σ ∈ (0.04, 0.1) where the steady state is achieved. We see reasonable agreement with the expected scaling relationship.

### 6.5 Probability of diversification as a function of initial community size

**Figure 16:**
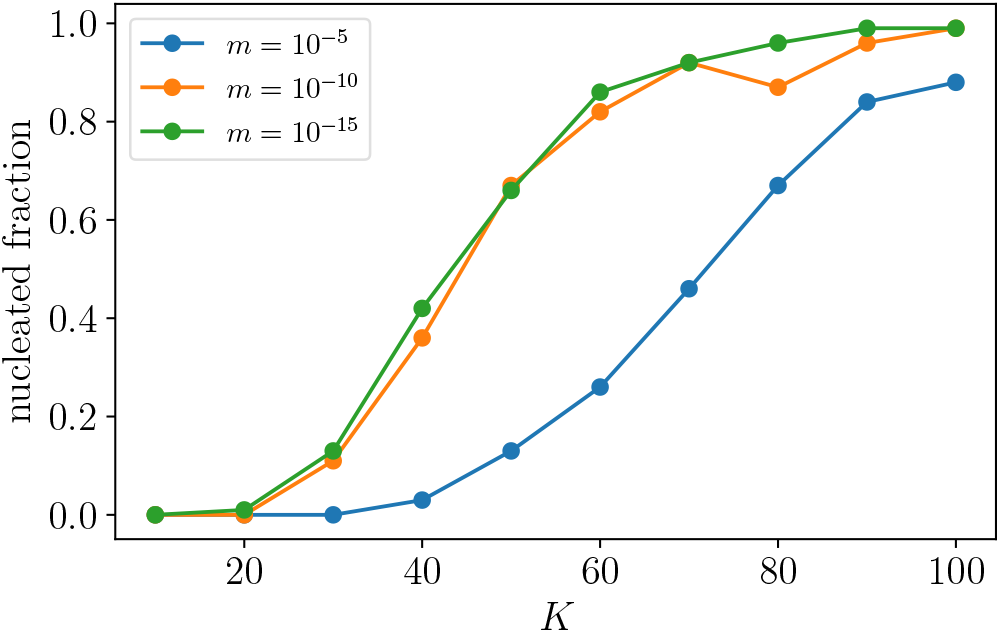
The probability of establishing the steadily diversifying regime increases with the initial number of strains and decreases with the migration rate. Although changing the migration rate from 10^−5^ to 10^−10^ changes the establishment probability significantly, further reducing *m* has little noticeable effect, indicating that the dependence of establishment probability on *m* is rather weak over several orders of magnitude. Data are averaged over 100 simulations for each set of parameter values. If *m* = 0, then migration cannot stabilize the spatiotemporal chaos, and so the establishment probability should vanish — but this is a very singular limit. Similarly as *γ* → − 1 we expect that the *K* needed to establish the diversifying phase will diverge.

### 6.6 Number of persistent strains with exponential 𝒫(*s*) in perfectly antisymmetric model

The antisymmetric model (ASM) with *γ* = −1 and no migration (considering only a single island) was analyzed in PAF. There is a unique uninvadable fixed point characterized by abundances 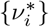 with half the strains having gone extinct. However the fixed point is marginally stable. Indeed, there is a family of chaotic steady states, parametrized a temperature like quantity *θ* which is roughly the range of the fluctuations in log *ν*_*i*_. However the average abundances are 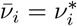.

Since there is no migration, the relationship between the biases and abundances is especially simple, as detailed below, the critical bias is *ξ*_*c*_ = 0 and the mean *s* is simply 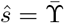. In the limit *K*Σ^2^ ≫ 1, the distribution of the drives with scale 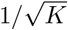 will be much narrower than that of the *s*’s. Therefore the scale of the *x*_*i*_ ≡ *ζ*_*i*_ + *s*_*i*_ distribution, *p*(*x*), will be Σ and it will decay exponentially for large *x*. We can get the distribution of *x* from the convolution of an exponential and gaussian distribution, with the drives having variance 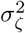:

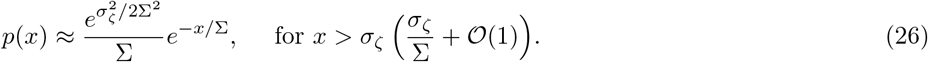

With the Lagrange multiplier 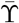, the average abundances 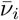 at the mean field fixed point will be 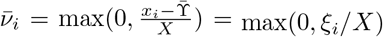 with *X* the total static susceptibility. Let us define *ϕ* as the fraction of strains that persists with positive bias. When 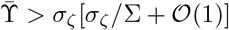, we can make the approximation

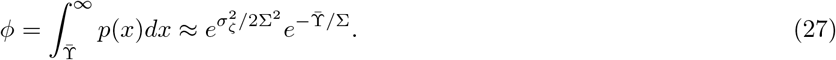

Self consistency requires that the average abundance over the initial *K* strains is

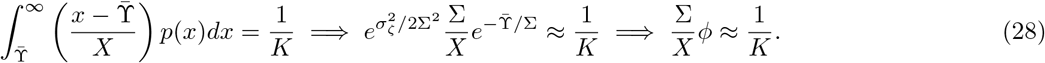

Now we combine this relation with the self consistency condition

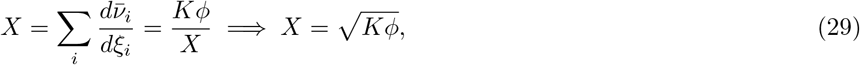

to obtain 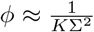. Therefore the number of persistent strains, when *K*Σ^2^ ≫ 1 is Σ^−2^, with coefficient one.

To obtain the behavior for arbitrary *K*, one must solve the mean field self consistency equations, with the exact distribution

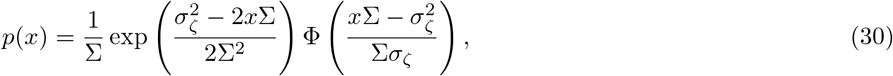

where Φ is the standard normal cumulative distribution function. There are three mean field equations for the three unknowns 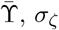 and *X*.

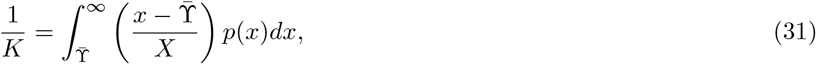

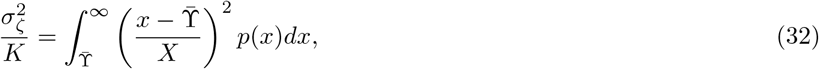

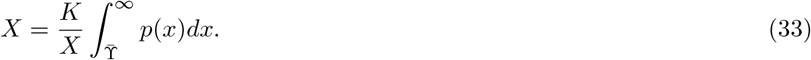

The number of persistent strains is 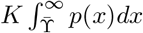, where 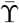 solves the above equations. Since for perfectly antisymmetric *V*, 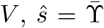 the solution of these allows us to check that, indeed, *ŝ*/Σ increases with Σ, as seen in Figure 5. Though one could solve the mean field equations numerically, we can obtain asymptotic results using the approximation in Equation 26.

For *K*Σ^2^ ≫ 1, we have 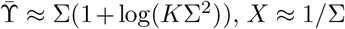 and 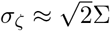, which is consistent with the asymptotic *Ansatz* for *p*(*x*) for the present strains: since 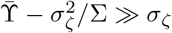 all persistent strains have large enough *x* for the approximation to be valid. Then the distribution of 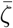 for the *persistent* strains is

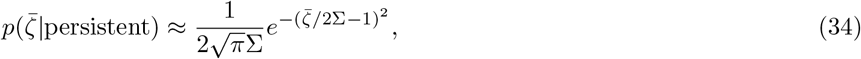

valid except for 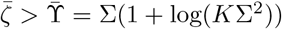 which is a negligible fraction of the distribution. The width of both the 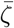 and the *s* distributions for the persistent strains is of order Σ. As expected more generally, these are comparable once *ŝ* is in the tail of the *s* distribution.

### 6.7 Estimation of drive and bias

Here we use the notation 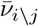 for the average abundance of strain *i* in the absence of strain *j*. The averaged force on a strain (excluding migration) can be measured in simulations. In the mean field approximation, this is decomposed as the sum of its bias and feedback from it perturbing the other strains::

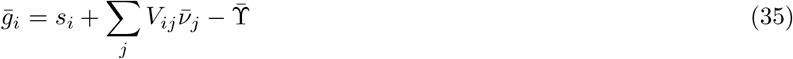

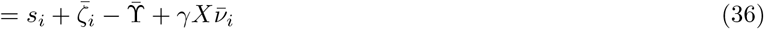

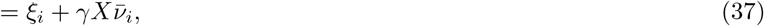

where the drive, 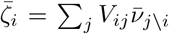, can be written in terms of mean field quantities as 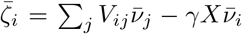 The term 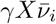 captures both the direct interaction and the perturbations to 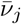 from strain *i*. Therefore, if we know *X*, we can calculate the drive and bias of a strain, since 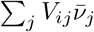 and 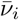 are measurable in our simulation, so we need only subtract off the 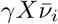 term from the average force due to the other strains. We can find *X* using the self consistency condition that 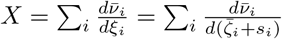

First, we make a guess for the susceptibility *X* which allows a provisional calculation of the 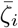. Then, using our numerical simulation data, we fit parameters *a, b* and *c* to a functional form

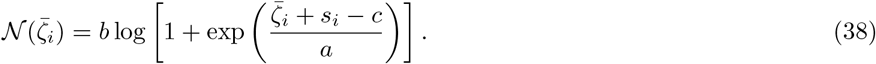

giving the mean abundance in terms of the drive plus *s*.

The justification for this form is, first, that it gives 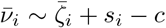 for 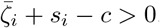. This is the expectation that 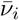 becomes proportional to *ξ*_*i*_ for positive *ξ*_*i*_ — though here we include an offset via the parameter *c* which roughly represents the expected correction to the linear function 𝒩(*ξ*) at large bias. The quantity 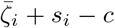 is similar to *ξ*_*i*_ but not quite the same due to a systematic difference between *c* and 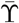. For large negative bias, the fitted form captures the roughly exponential decrease of 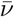 due to the rareness of blooms.

The data are fit by this functional form quite well. We observe that *c* tends to be smaller than the measured 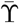 within a given epoch, which also could be due to the effect of migration which elevates 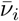 for strains with negative *ξ*_*i*_. The parameter combination *b*/*a* is roughly equal to −1/*γX*, as expected from the mean field analysis. From the hypothesized form of 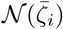 we can recalculate the susceptibility via 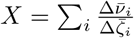 (where the derivative is calculated numerically). We then update our value of *X* by averaging the old guess with the new estimate.

By iterating over the susceptibility until it converges to a value where the assumed drives 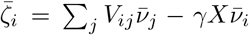 reproduce the susceptibility via the self-consistency condition, we can get an estimate for the susceptibility and therefore the drive as well. Then the bias is 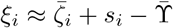, where we have subtracted the Lagrange multiplier (neglecting effects of order 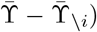 and added back in the nonspecific fitness part. We can validate that our bias estimator is reasonably accurate by comparing the inferred biases to the biases of strains in their first epoch of invasion — the latter can be measured directly from simulations since they are simply the invasion eigenvalues for incoming strains. This works well, so we extend it to infer the biases for all the other strains in the community.

In contrast to the assembled communities studied in PAF, we cannot use the condition that the drive averaged over all the initial *K* strains, is zero. PAF used this — along with the expected truncated gaussian shape with variance 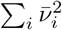 and lower limit 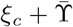 — as a check in calculating the drives. However conditioning on evolution means that the mean drive is no longer 0 and the drive distribution is no longer a truncated gaussian, so we must rely on the method described above. To increase the quantity of data on which to perform our fit for 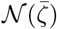 in a given epoch, we pool data of strain abundances and forces from up to 20 epochs around this focal epoch, provided their *L* is within 5% of the *L* in the focal epoch, with the idea that these communities are statistically similar and therefore have a similar 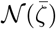.

The function 𝒩(*ξ*) for the evolved communities with *ρ* = 0 looks quite similar to the function for assembled communities However for communities evolved with *ρ* = 0.95, 𝒩(*ξ*) looks substantially different. This indicates that the build-up of correlations in the community requires modifications of the independence assumptions of the mean field theory for the assembled community.

**Figure 17:**
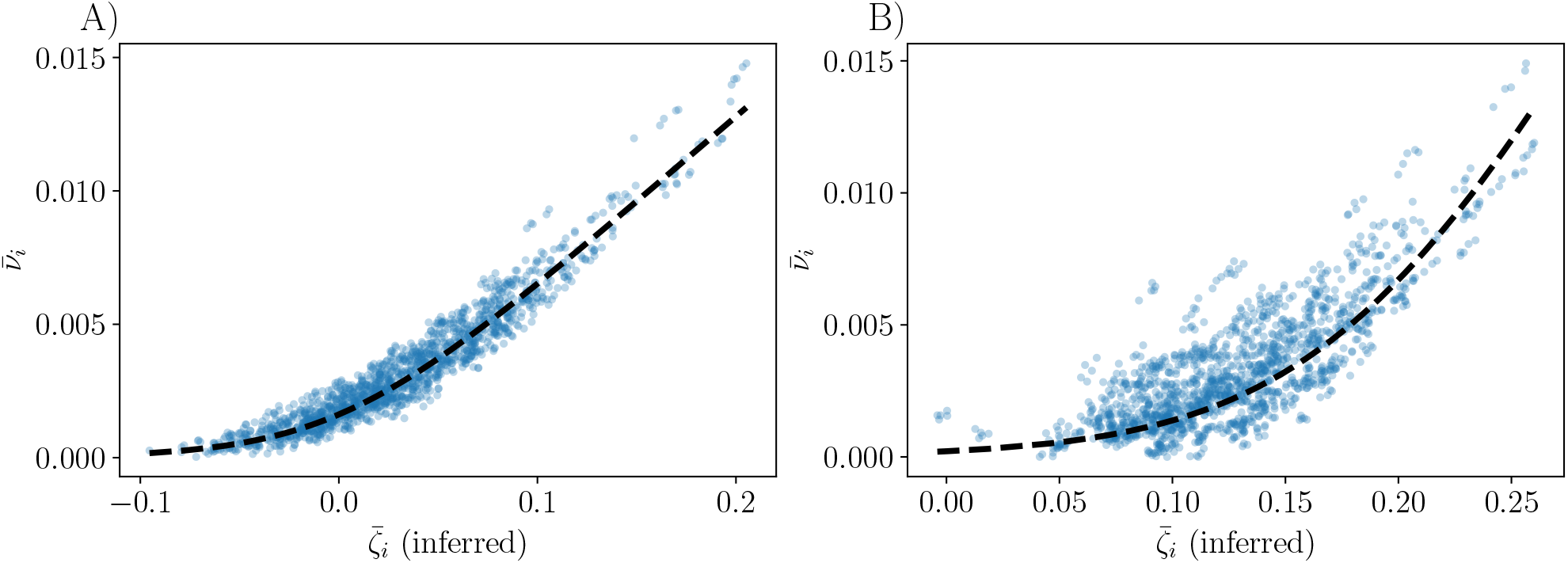
Inference of the drive for (A) communities evolving with *ρ* = 0 and (B) communities evolving with *ρ* = 0.95. In both cases, there are no nonspecific differences: all *s*_*i*_ = 0. The black lines shows the fit of the functional form for 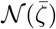 after *X* has converged to a self consistent value. Data are pooled (both for fitting and for plotting) over 5 consecutive epochs, each with ~ 300 extant strains. For *ρ* = 0.95 it seems that correlations in the interactions that have accumulated due to the relatedness is affecting the relationship between 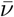 and 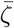. Certain strains are visible outliers from the average dependence of 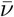 on 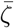, and appear multiple times on plots since data are pooled across 5 epochs.

From the inferred function 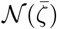, and the measured mean abundance of every strain in the community, we can calculate the fragility, defined as 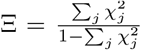. This closer 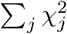 is to 1, the more unstable the community is to perturbations. Indeed we find that for a simulations diversifying from 50 to 500 strains, 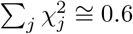 for the evolved community of ≅ 500 strains, while early in the simulations, when *L* < 100, we find 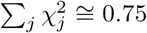. When comparing 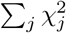 for assembled and evolved communities, both of 500 strains, there is not a clear difference.

### 6.8 Scaling solution for Fokker-Planck equation without general fitness differences

Without general fitness differences or correlations, the evolution of the average drive of a strain in the Markovian approximation obeys a Langevin equation (Equation 10). With a source term corresponding to incoming strains, this yields a Fokker Planck equation (Equation 11) for the number density 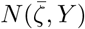 whose integral at any time gives us the number of strains. The crucial *Ansatz* is that all the quantities (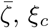, *ξ*_*c*_ and 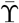) scale as 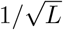, and we now take *L* to be a function of *Y*. Then we can hypothesize a scaling solution 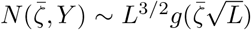 for some function *g* so that 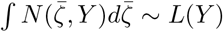, as the integral over the distribution of drives gives us the total number of strains. For ease of notation we define the similarity variable 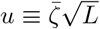, and derivatives with respect to *u* by primes: then Equation 11 becomes

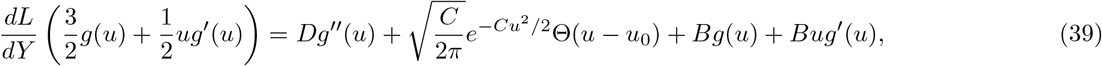

where we define 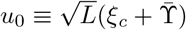. We can make an *Ansatz* of the form

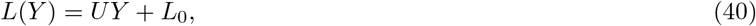

where the coefficient *U* is the average rate of increase or decrease of the diversity per invasion attempt.

The equation for *g*(*u*) then becomes

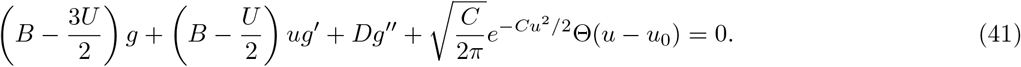

We can solve this equation numerically with boundary conditions *g*(*u*_0_) = 0 and *g*(∞) = 0 (exact solutions are available for certain values of *B, D* and *U*). The first of these boundary conditions comes from the extinction criterion that enforces 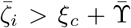 for every extant strain. The solution depends on these parameters in addition to *u*_0_. Then, by enforcing 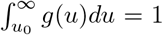, we can fix the value of *U*, which depends on *u*_0_, *B* and *D*. One can see, numerically, that for *B* and *D* both large, the consistent value of *U* is negative, implying loss of diversity.

In Figure 8, we use the measured values of *U* and *C*, and adjust the values of *B* and *D* to give a normalized function *g*(*u*).

### 6.9 Scaling solution for Fokker-Planck equation with independent general fitness differences

If incoming strains have independently drawn nonspecific differences, *s*_*i*_, then we can write an evolution equation for the joint distribution of the drives and *s*_*i*_’s

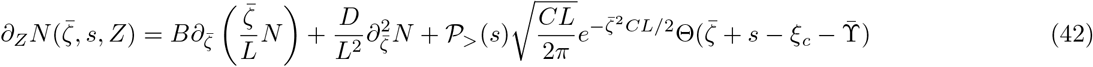

where we abbreviate 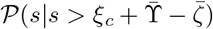 by 𝒫_>_(*s*). However, unlike the case with general fitness differences, now 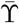 does not scale as 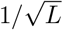. Indeed, the scaling of 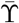 with *Z* determines how *L* depends on *Z*. If we specify the form of 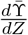, then we can look for solutions which travel up in the s direction at the same speed as 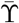.

We can define similarity variables 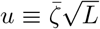 and 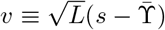. Then we look for solutions of the form 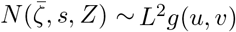 so that the integral of *N* over 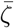 and *s* is proportional to *L*. The boundary condition is that *g* vanishes along the line *u* + *v* = *u*_crit_. Plugging this into our Fokker Planck equation, and defining 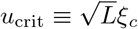, we obtain

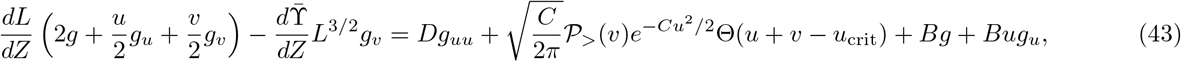

where subscripts denote derivatives of *g*. We can see that the 𝒫_>_(*s*) distribution generically breaks the scaling — because it depends on 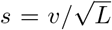, not only on *v*. But we can look for solutions which obey the scaling with 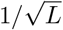. Take 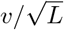 as exponentially distributed with scale Σ. Then in order for our scaling *Ansatz* to be valid, the quantity 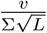 must be independent of *L*, so we must have 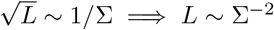. Then this requires 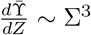 in order to make the left hand side of Equation 43 independent of *L*. Furthermore we have *dL*/*dZ* → 0 since *L* ~ Σ^−2^. Therefore we have

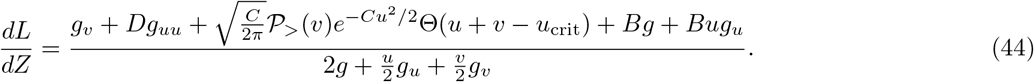

### 6.10 Joint distribution of drive and *s* of invader

Since the distribution of the invader 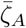 and *s*_*A*_ are independent, we can write their joint distribution as

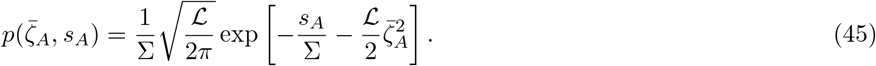

Conditioning on successful invasion adds a factor of 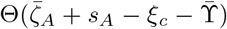, which enforces that *ξ*_*A*_ > *ξ*_*c*_. This would also change the normalization constant of the distribution — so we define a constant *J*, the reciprocal of the invasion probability, such that 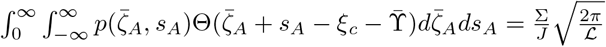. We can then calculate the marginal distribution of *s*_*A*_ conditioned on invasion:

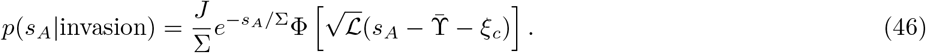

Similarly, the distribution of 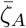 conditioned on invasion is

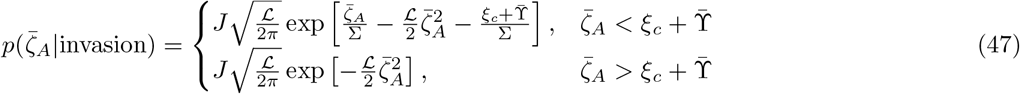

We can also calculate the distribution of 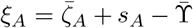 conditioned on invasion. In the limit that 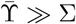 we can take the *s* integral over both positive and negative *s* since the negative part contributes negligibly. One then obtains

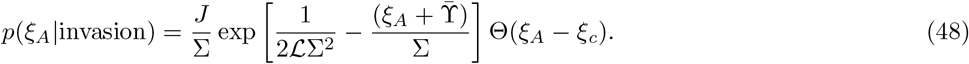

One can carry out by simple extension the same calculation if the *s*’s are correlated but 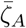 is still gaussian distributed with mean 0 and variance ℒ (i.e. the interactions in *V* are uncorrelated), but we have not done so here.

### 6.11 Adaptation and diversification rates versus attempted invasions

One signature of evolution with nonspecific differences is that invasion by an uncorrelated strain becomes more difficult as the community matures, due to an increase in ŝ and therefore 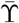. We thus expect that the number of strains and average general fitness of the community, ŝ, increase as the log of the number of attempted invasions raised to a power determined by the scaling analysis from Section 4.5.

**Figure 18:**
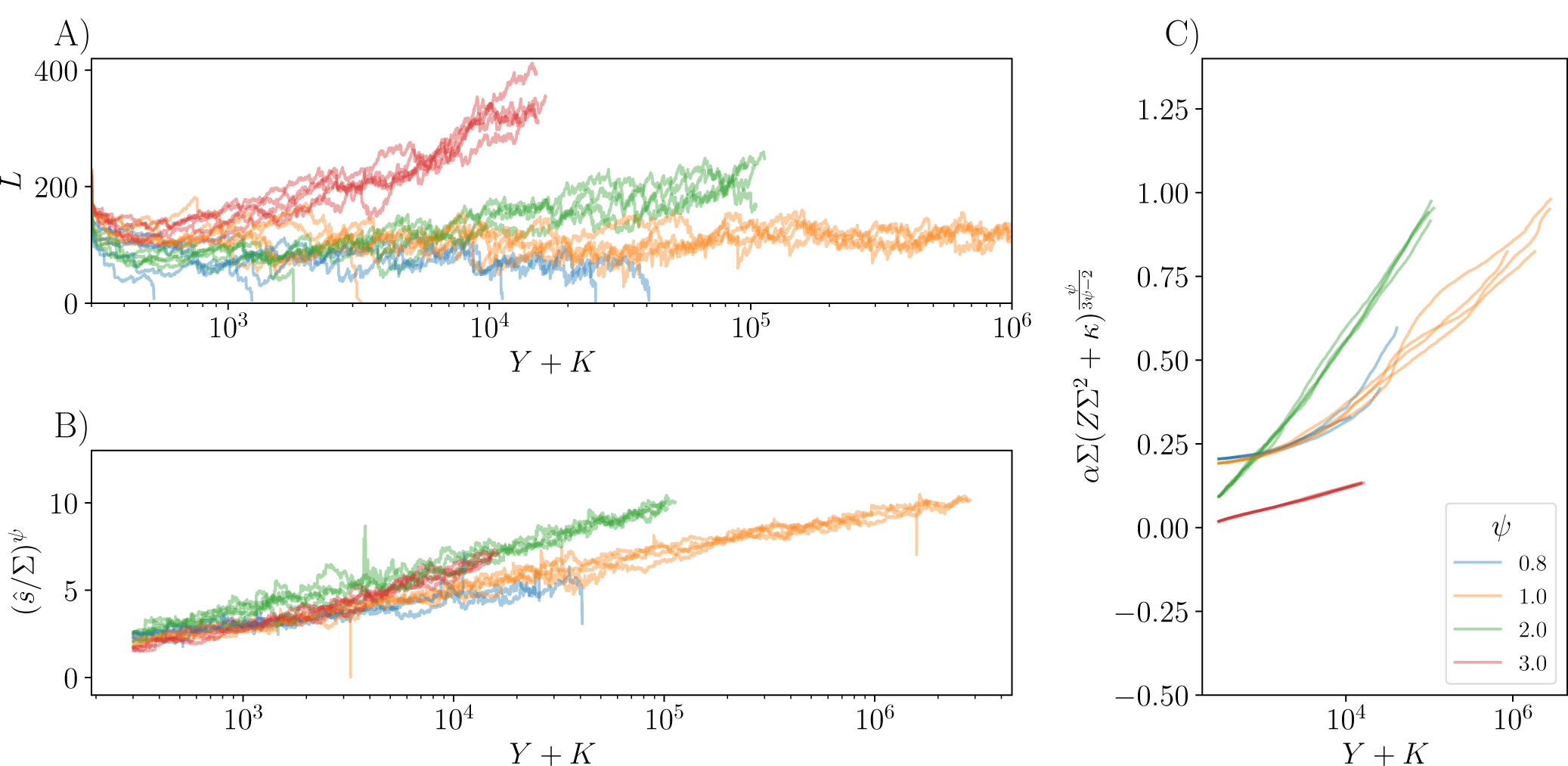
Theory yields predictions of relationships between the rate of successful invasions as a function of attempted invasions. (A) The number of strains versus number of attempted invasions *Y*, with offset *K*. (B) The mean general fitness versus attempted invasions. With these rescaled axes, theory from Section 4.5 predicts that the curves should be straight lines with similar slope. (C) The relationship between ŝ and *Z*, in conjunction with the relationship between ŝ and *Y*, allows one to relate the number of successful and attempted invasions (*Z* and *Y*) directly. On this plot we expect that all curves have the same slope once they have reached the asymptotic regime where 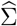 is changing slowly. However it is likely that for *ψ* = 3, we have not yet reached that regime. As in the data from Figure 9, Σ = 0.066, 0.08, 0.176, 0.173 for *ψ* = 0.8, 1, 2, 3 respectively.

The fitting parameters *α, β* and *κ* used in Figure 9 (for each value of *ψ*) are determined by trial and error, and listed below.

| $\psi$ | $\Sigma$ | $\alpha$ | $\beta$ | $\kappa$ |
| --- | --- | --- | --- | --- |
| 0.8 | 0.066 | 0.0028 | 6.78 | 15.72 |
| 1 | 0.08 | 0.16 | 0.61 | 15 |
| 2 | 0.176 | 0.92 | 0.59 | 3 |
| 3 | 0.173 | 0.96 | 0.81 | 1.92 |

### 6.12 Statistics of *V* conditioned on evolution

To check whether the evolved interaction matrices, studied in Section 4.8, are statistically significantly different from the matrices of the pool, we normalize each statistic of the evolved interaction matrices by the standard deviation of the estimator of the corresponding statistic, from matrices drawn from the original pool without any conditioning. This gives us an idea of how atypical the measured statistics are for the matrices of size *L*× *L* with *L* = 500. The quantities we have analyzed are where Stde 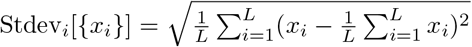 (the population standard deviation) and Corr[*X, Y*] is the Pearson correlation coefficient calculated for a sample (the covariance normalized by the standard deviations).

By subtracting off the mean of each quantity and dividing by its standard deviation in the pool, we obtain a measure of how significant the changes due to evolution are. Therefore we define quantities 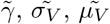 and 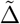 to be the empirical matrix statistics measured in units of standard deviations of the pool away from the pool mean.

Although a considerable number of statistics are significantly different from their values without any conditioning, few incur substantial changes, with the largest change by far occurring in the *µ*_*V*_ for the simulations with *ρ* = 0.95; however even this is a relatively small effect.

### 6.13 Replacement probability of a parent by a mutant: Lotka-Volterra approximation ignoring migration

In this section we analyze the effective Lotka Volterra equations emerging from correlated dynamics of mutant and parent, as introduced in Section 4.6. This analysis is appropriate only for strains with positive bias — and even then approximate — since we neglect the effects of migration and average the dynamics of log *ν* to obtain equations for parent and mutant frequencies *ν*_*P*_ and *ν*_*M*_ :

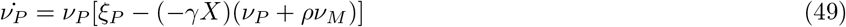

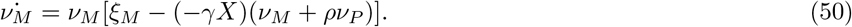

Numerics indicate that dynamical correlations between *ν*_*P*_ and *ν*_*M*_ make it much easier for closely related strains to coexist than in simple stable dynamics. The following results provide a null model against which one can compare the dynamics of parent-mutant replacement in the evolving STC.

In the absence of the mutant, 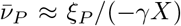 (which is the correct behavior for large positive *ξ*_*P*_). The condition for invasion of a mutant into an environment containing only the parent is that its invasion eigenvalue, suppressed by the effects of the parent, is greater than the critical bias which amounts to *ξ*_*M*_ − *ρξ*_*P*_ > *ξ*_*c*_ and the condition for invasion of the parent into the mutant is likewise *ξ*_*P*_ − *ρξ*_*M*_ > *ξ*_*c*_. In terms of the drives, these conditions are 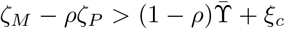 for the mutant to invade the parent, and the same condition with *M* and *P* flipped for the parent to invade the mutant. With the mutant drive parameterized as 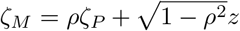 with *z* a gaussian random variable with mean 0 and scale *ζ*_*P*_, the probabilities of coexistence between parent and mutant, and of replacement of the parent, are

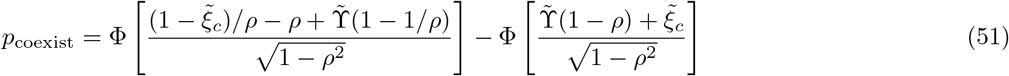

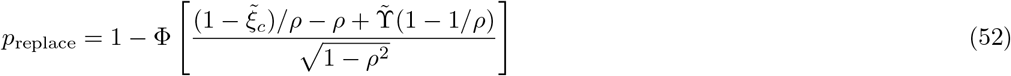

where Φ is the standard normal cdf, and 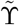 and 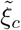 are the Lagrange multiplier and critical bias, both normalized by the scale of the *ζ* distribution. Note that the expressions are only valid for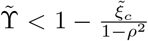 : otherwise, in this approximation, the coexistence probability vanishes and all invasions are replacements. The probability of mutant invasion 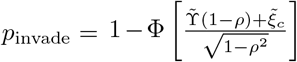. For 1 − *ρ ≪* 1 and 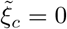, the invasion probability is 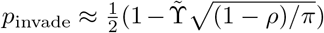 and the coexistence probability predicted by this simple approximation is 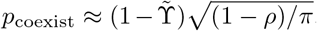. With *ρ* = 0.99 and 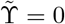 (in the STC 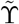 should be nonnegative, and setting it to 0 maximizes the coexistence probability), this predicts that the probability of coexistence *conditional on the mutant invading* is *p*_coexist_/*p*_invade_ ≅ 0.12 which is much smaller than the observed *p*_coexist|invade_ > 0.5 (Figure 10).

The main problem with this approximation is that it needs to be used when the biases are close to the critical *ξ*_*c*_ which is substantially negative, while the approximation of ignoring migration certainly breaks down in this regime. Properly understanding the coexistence of replacement of related mutant and parent requires a fuller understanding of their coupled dynamics, including migration.

